# Deep DNAshape: Predicting DNA shape considering extended flanking regions using a deep learning method

**DOI:** 10.1101/2023.10.22.563383

**Authors:** Jinsen Li, Tsu-Pei Chiu, Remo Rohs

## Abstract

Understanding the mechanisms of protein-DNA binding is critical in comprehending gene regulation. Three-dimensional DNA shape plays a key role in these mechanisms. In this study, we present a deep learning-based method, Deep DNAshape, that fundamentally changes the current *k*-mer based high-throughput prediction of DNA shape features by accurately accounting for the influence of extended flanking regions, without the need for extensive molecular simulations or structural biology experiments. By using the Deep DNAshape method, refined DNA shape features can be predicted for any length and number of DNA sequences in a high-throughput manner, providing a deeper understanding of the effects of flanking regions on DNA shape in a target region of a sequence. Deep DNAshape method provides access to the influence of distant flanking regions on a region of interest. Our findings reveal that DNA shape readout mechanisms of a core target are quantitatively affected by flanking regions, including extended flanking regions, providing valuable insights into the detailed structural readout mechanisms of protein-DNA binding. Furthermore, when incorporated in machine learning models, the features generated by Deep DNAshape improve the model prediction accuracy. Collectively, Deep DNAshape can serve as a versatile and powerful tool for diverse DNA structure-related studies.

## Main

Binding interactions between DNA binding proteins such as transcription factors (TFs) and their DNA target sites are crucial for gene regulation; thus, it is vital to fully understand TF-DNA binding mechanisms. These mechanisms can be viewed from two perspectives: base readout and shape readout^1,2^. Base readout occurs through direct contacts between amino acids and DNA bases, and a relatively strict pattern is followed. For instance, the zinc finger TF family heavily relies on base readout to recognize DNA targets^3,4^. Shape readout happens when proteins interact with the double helical shape, rather than interacting directly with certain atoms through hydrogen bonds on nucleobases. As an example, proteins containing positively charged amino acids (e.g., arginine, protonated histidine, and lysine) may favor a certain DNA shape, such as a narrower minor groove that is more negatively charged^1^. Some TFs from the homeodomain family utilize shape readout in specific DNA regions within the core binding site^5–9^. Narrower minor groove regions can result from diverse DNA sequences, providing an additional signal for binding specificity. Another example is the myocyte enhancer factor-2 (MEF2) family, which uses shape readout in the center of its degenerate motif^10^. In general, TFs use a combination of base and shape readout modes to achieve DNA binding specificity. The mismatch-induced DNA shape alterations that influence TF-DNA binding affinity prove the importance of shape readout^11^. However, the influence of the base and shape readout mechanisms can be difficult to parse.

The influence of shape readout in DNA binding mechanisms extends to various aspects of DNA structural features, or DNA shape. To measure shape readout quantitatively, one must first define DNA shape. This article focuses on the B-form representation of DNA molecules - the canonical right-handed double helix. Three-dimensional (3D) DNA shape describes the global and local relationships between nucleobases, base pairs, and sugar-phosphate backbones^12^. Although DNA oligonucleotides are flexible molecules with varying DNA shape features, certain optimal conformations may be intrinsically favorable, as seen in their static or sometimes equivalent time-averaged representation of DNA shape features^13^. Proteins that favor certain intrinsic DNA shape features may bind DNA targets with other DNA shape; however, they may require a higher energy to maintain the binding, thus exhibiting a lower binding affinity ^6^.

The energy required to alter an unfavored static DNA shape may depend on the DNA flexibility. Therefore, it is important to consider the conformational flexibility of the DNA and fluctuations in its shape features. As revealed by, for instance, molecular dynamics (MD) simulations, DNA shape features fluctuate in various ways^14–16^. For example, CpG, CpA and TpA base-pair (bp) steps are generally more flexible^17^, whereas A-tracts are conformationally rigid^17,18^. The length of A-tracts has also been observed to impact the flexibility of neighboring regions^19^. Methylated cytosine induces DNA flexibility changes that influence nucleosome stability^20,21^. Another recent study using MD simulations showed that conformational flexibility contributed by the flanks plays an important role in homeodomain binding^22^. However, no high-throughput methods have been proposed yet to predict DNA shape fluctuations.

Methods such as MD or Monte-Carlo (MC) simulations and X-ray crystallography (XRC) can be used to acquire DNA structures for short DNA fragments. Methods such as Curves^12,23,24^ and 3DNA^25^ have been developed to derive DNA shape features from computational trajectories or experimentally solved structures. High-throughput methods such as DNAshape^26–28^ have been introduced to circumvent the difficult and sometimes impossible task of simulating or experimentally solving structures. Numerous studies have successfully employed the DNAshape method ^26^, demonstrating the effectiveness of using high-throughput methods to predict DNA shape features^29–33^.

Nevertheless, although DNAshape^26–28^ is an efficient method, it relies entirely on a pentamer query table containing all possible pentamers compiled from extensive MC simulations^34^. The pentamer length limits this method because only the nearest and next-nearest neighbors are accounted for when considering the influence of the sequence environment on the center of the pentamer. As for other data sources, such as MD simulation data and experimental structures in the PDB, only tetramer query tables could be generated due to data paucity^27^. Therefore, effects from longer-range neighbors are totally neglected in this query table setup.

Our new approach, Deep DNAshape, overcomes the limitation of DNAshape, particularly its reliance on the query table search key. This advancement is pivotal, given that the limitation was only caused by the available amount of data. Deep DNAshape enhances the capability to discern how the shape at the center of a pentamer region is influenced by its extended flanking regions, providing a model that offers a more accurate representation of DNA.

The pentamer query table contains all possible pentamers for DNA shape features of the central base pair, considering up to a 2-bp flanking region, which consists of the nearest-neighbor and next-nearest-neighbor. Despite this definition, flanking regions exceeding two bp may still be influential, which has been shown for the central TpA step^35^. For some TFs having a long core motif, a more complete view of DNA shape considering longer-range flanking regions is necessary. In addition, some DNA shape values are bimodal^15^, and statistically calculating an average value for the query table may not capture the whole picture. The ability to approximate DNA shape features using only sequence information such as mono- and di-nucleotides^36^ also highlights the limitations of the pentamer query table. Although it is possible to generate a query table for longer *k*-mers, the number of simulations that would be necessary to cover all longer *k*-mers is exponentially higher; meanwhile, there are not enough existing experimental structures^27^. Therefore, there is a need for a method that can accurately predict DNA shape features in a high-throughput manner using only limited data while considering longer-range effects.

To develop such a model, we began with assumptions about how the 3D DNA structure is affected by sequence. Firstly, we assumed that DNA shape features at each base pair are mainly influenced by their neighboring base pairs, and that this influence weakens with distance. Secondly, we assumed that this influence can be statistically inferred. Therefore, we designed a specialized deep learning architecture to deal with variable-length DNA sequences and computed the neighboring effects of flanking regions in a layer-by-layer manner (Figs. 1a-d, S1). We trained the model on DNA shape features that were previously analyzed and compiled from MC simulations (Fig. 1b, S2-3) and which had been experimentally validated^26^. The model now considers longer range neighboring effects compared to current data source limitations (Fig. 1e) and can be used for predicting DNA shape features from any given sequence (Figs. 1f-g). We then evaluated the predicted DNA shape features with a tetramer query table derived from MD simulations (Table S1)^27^. We compared predictions from our resulting model, Deep DNAshape, to predictions from our original pentamer-based DNAshape (DNAshapeR) method^37^ (Figs. 2, 3, 4, Table S2). To thoroughly benchmark the model, we also trained it on alternative DNA shape sourced from experimentally solved structures^38^ and MD simulations^39^ (see Supplementary Information), leading to the creation of Deep DNAshape (Expt) and Deep DNAshape (MD). These models were then compared with the MC-trained Deep DNAshape, along with comparative analysis across all model variants (Fig. S4, Table S3).

**Fig. 1.**
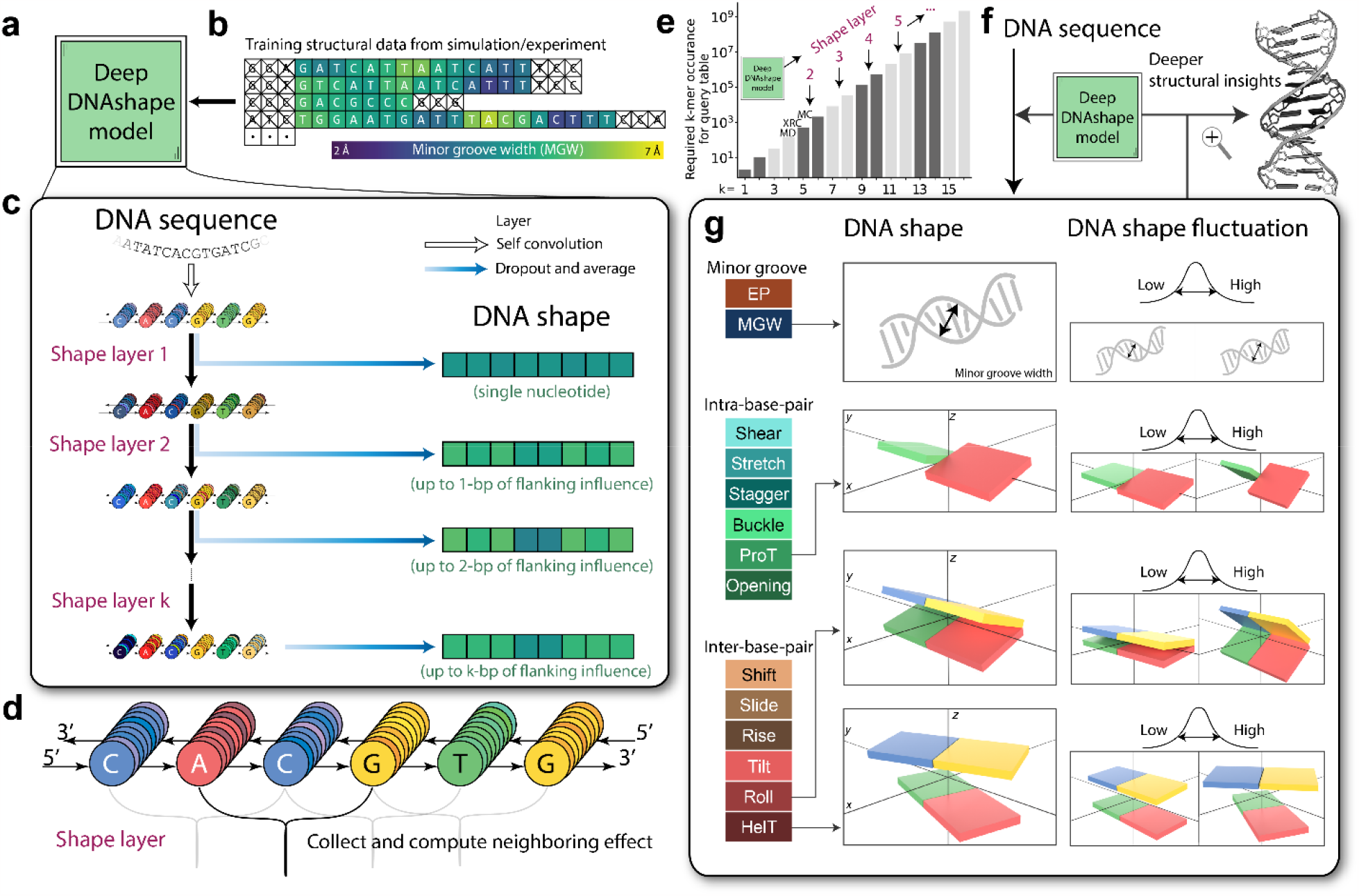
Deep DNAshape schematic. **a, b)** Deep DNAshape model (a) and training data for the model (b). Sequences used in simulations are pooled together, and corresponding DNA shape values are assigned to each position of each sequence. Some values, such as MGW at the terminal base pairs, are not defined and therefore excluded. **c)** Schematic representation of the Deep DNAshape model. DNA sequences go through several ‘shape layers’. Each time the model goes through a shape layer, features for each individual position are passed to its nearby two positions, allowing one additional consideration for the flanking region. **d)** Simple diagram of ‘shape layer’. In each layer, features from neighboring nodes are collected and computed with information in each current node. Features are updated for each node on the sequence. See Methods for details. **e)** Capacity comparison between the Deep DNAshape model and current data limitations. The shown labels “MC”, “XRC” and “MD” are current limitations to generate pentamer or tetramer query tables sourced from Monte Carlo simulations, X-ray crystallography, and molecular dynamics simulations, respectively. **f)** Deep DNAshape model usage. Deep DNAshape can process a given DNA sequence as a string of characters (A, C, G and T) and predict any specific DNA shape for each nucleotide position of a sequence. **g)** All DNA shape predicted by Deep DNAshape model. In addition to static DNA shapes, the model can predict DNA shape fluctuations (Cartoons do not represent real values of low or high. Values can be negative or positive for angle shape parameters.). Shown are graphical explanations of the four most-used DNA shape features (MGW, ProT, Roll, and HelT).

**Fig. 2.**
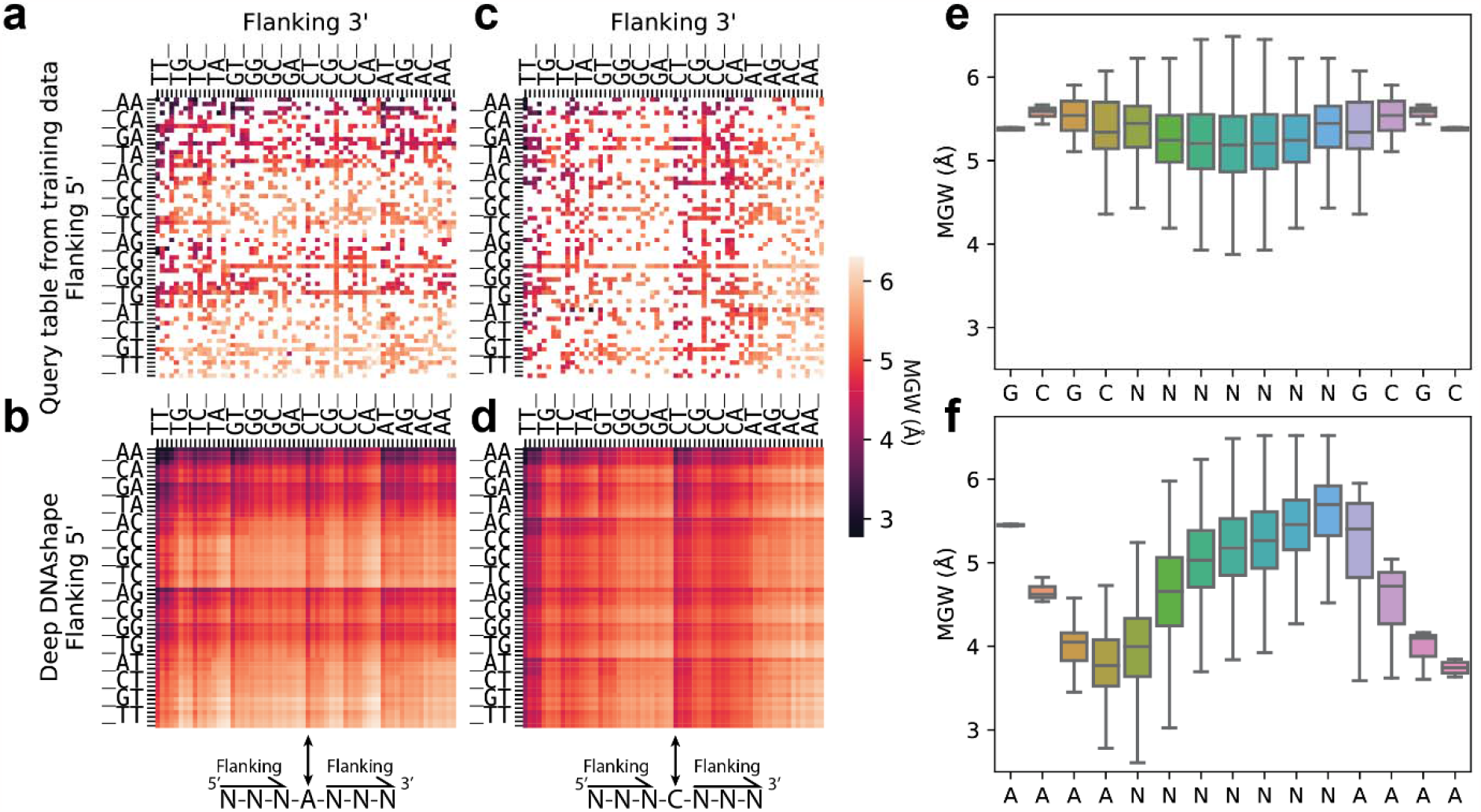
Minor groove width predicted by Deep DNAshape using extended flanking regions. **a-d)** Heatmaps showing MGW values of central position for all possible 7-mers. a) and c) show MGW values as if we constructed a 7-mer query table from all available MC simulations directly. b) and d) MGW predictions generated by the Deep DNAshape method for all possible 7-mers. a) and b) All sequences with ‘A’ base in the middle. c) and d) All sequences with ‘C’ base in the middle. Flanking regions from 5’ and 3’ are sorted based on distance to the center and base-pair character in the heatmap. ‘_’ represents A, C, G, and T in sequential order for flanking 5’ and in reverse for flanking 3’. **e-f)** Boxplots showing MGW values predicted by Deep DNAshape for random sequences with fixed 5’ and 3’ caps. e) Predictions are capped by ‘GCGC’, and f) Predictions are capped by ‘AAAA’. Center line indicates the median. Box limits are 75th and 25th percentiles. The whiskers extend 1.5 times the IQR from the top and bottom of the box. Outliers are removed in boxplots.

**Fig. 3.**
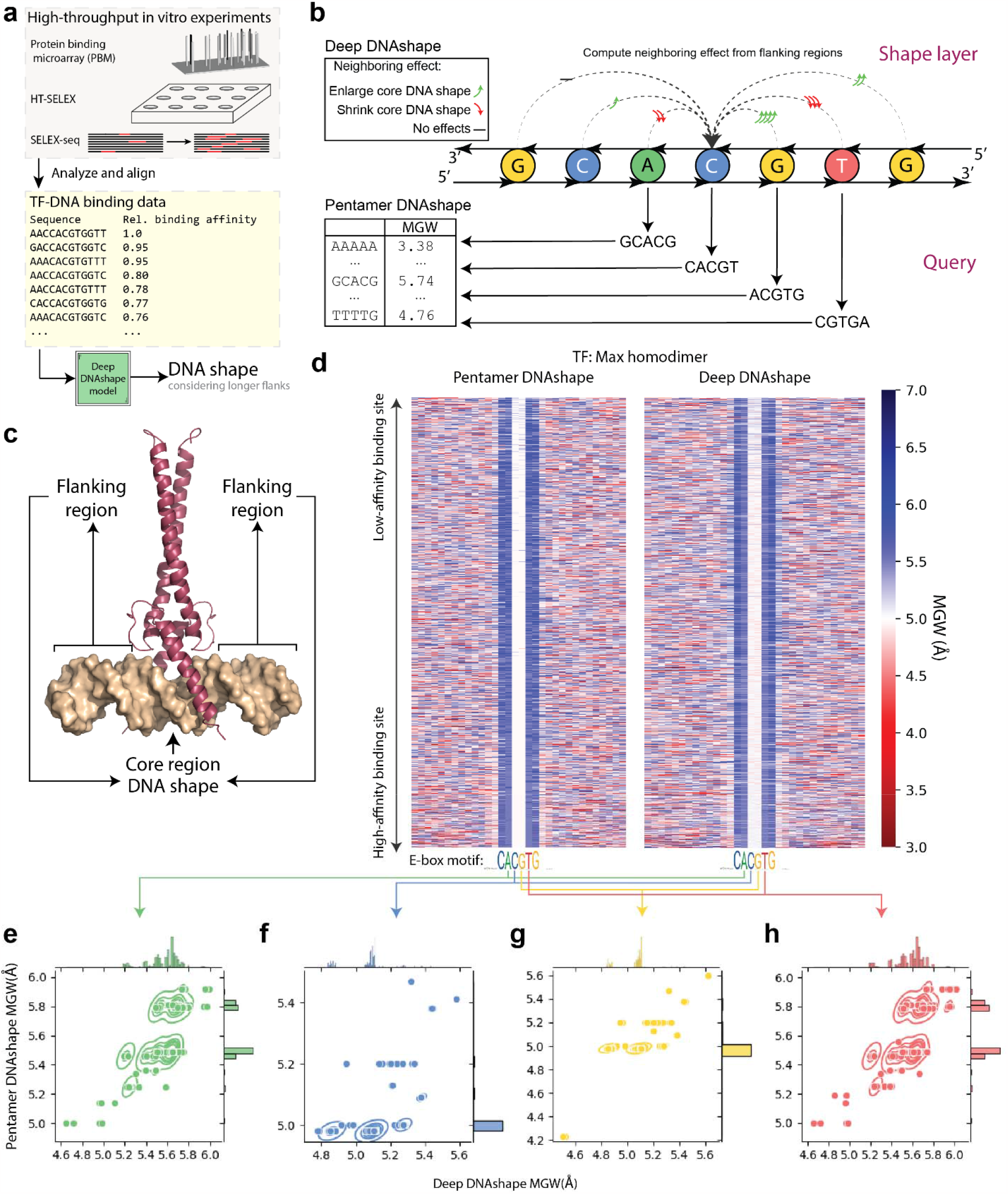
Comparison of Deep DNAshape and pentamer DNAshape methods on TF-DNA binding data. **a)** General pipeline to apply Deep DNAshape on TF-DNA binding assay data. **b)** Comparison of methodology to predict DNA shape features from Deep DNAshape and pentamer-based DNAshape methods. **c)** Co-crystal structure of DNA-bound Max protein homodimer (PDB ID: 1AN2). Flanking regions indicate regions without protein contacts. Core is the region with protein contacts. **d)** MGW predicted by original pentamer DNAshape vs Deep DNAshape for Max protein and DNA binding data, in order of relative binding affinity. Color represents DNA shape values. Data are aligned with core binding site. Only top 25% of binding data are used. **e-h)** Scatterplots of central four core MGW values predicted by Deep DNAshape and pentamer DNAshape. Gaussian kernel density estimation plot is added to show the contour of scatter density. Histograms along axes are shown to visualize the distribution of DNA shape values predicted by both methods. Values predicted by Deep DNAshape reveal structural variations in the center of the E-box that the pentamer-based DNAshape method was unable to expose.

**Fig. 4.**
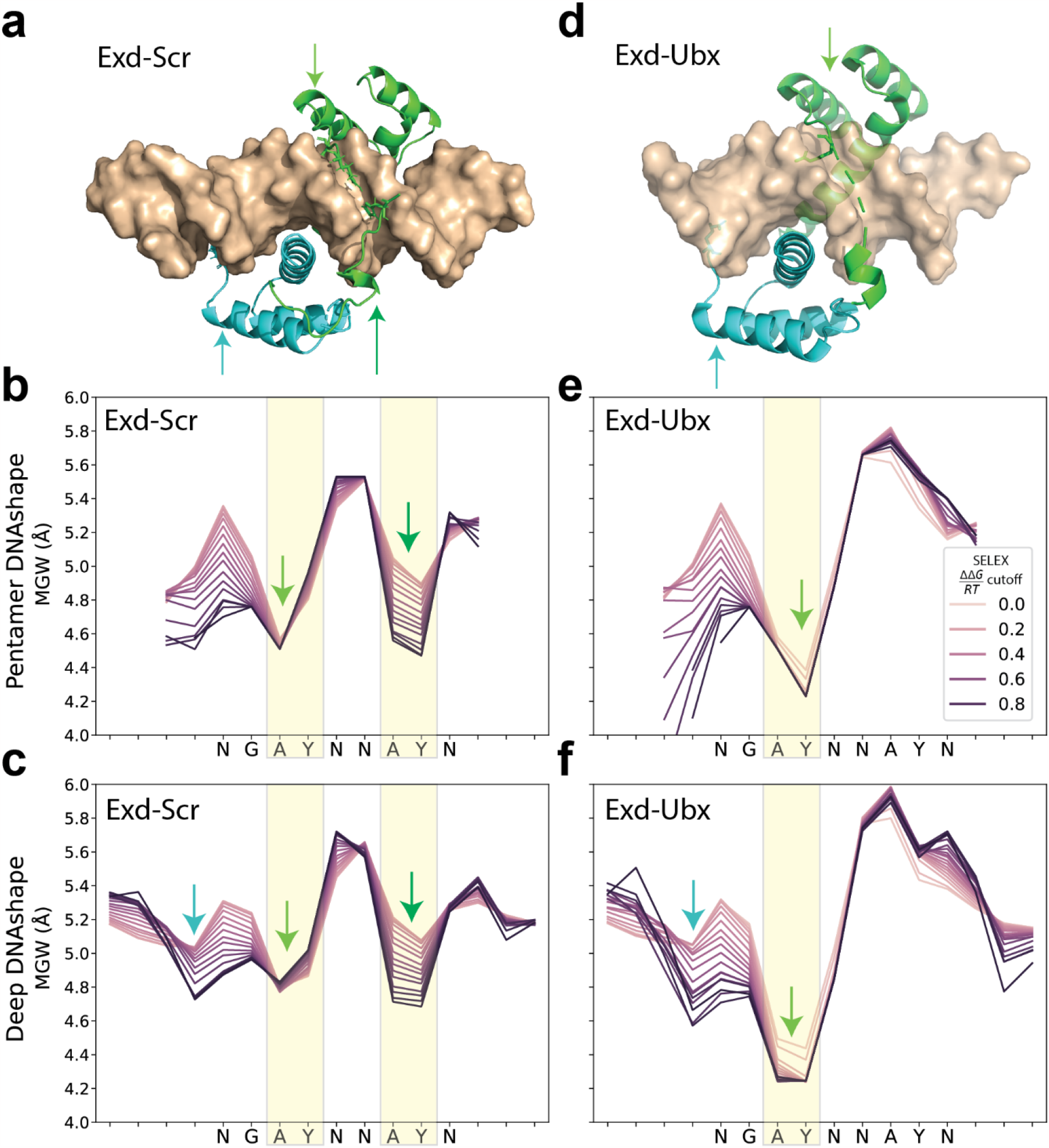
Evaluations of Deep DNAshape on extended flanking regions using Hox-TF binding data. **a)** Co-crystal structure of DNA-bound Exd-Scr heterodimer (PDB ID: 2r5z). Arrow indicates location of insertion from Exd (cyan) and Scr (green) residues into the minor groove of DNA. **b-c)** MGW predicted by original pentamer DNAshape and Deep DNAshape, for Exd-Scr SELEX-seq data. Lines are calculated based on cutoff values of relative binding affinities. Darker colors in plot correspond to sequences with higher binding affinities. The comparison of the two panels shows that Deep DNAshape predicts the MGW of the flanking regions and Exd minimum that the pentamer-based DNAshape method was unable to predict. **d)** Co-crystal structure of DNA-bound Exd-Ubx (PDB ID: 4cyc). Arrow indicates location of insertion from Exd (cyan) and Ubx (green) residues into the minor groove of DNA. **e-f)** MGW predicted by original pentamer DNAshape and Deep DNAshape, for Exd-Ubx SELEX-seq data. Lines are calculated based on cutoff values of relative binding affinities. Darker colors in plot correspond to sequences with higher binding affinities. The comparison of the two panels shows that Deep DNAshape predicts the MGW of the flanking regions and Exd minimum that the pentamer-based DNAshape method was unable to predict.

In addition, the design of the model unlocks the potential to examine systematically how DNA shape fluctuations are affected by extended flanking regions. We previously generated a query table containing standard deviation (SD) values for 13 DNA shape features and used them in a machine learning study^27^. Although these values were statistically computed, nevertheless, the model performance was significantly improved when the values were included^27^. We assumed that these SD values were highly correlated with true fluctuation values. Therefore, although conformational flexibility of DNA is important^40^, it is a shape feature that is difficult to access and frequently overlooked in research. Here, we used the same approach as in predicting the static DNA shape to predict DNA shape fluctuation (FL) values: specifically, we directly calculated fluctuation from MC simulations using Curves^12^. We investigated whether these high-throughput-predicted fluctuation values aligned with previous findings on DNA flexibility^40^, and we considered the insights that these fluctuation values might provide.

Next, we compared the new Deep DNAshape model with the original pentamer DNAshape method. We tested our model on data from TF-DNA binding assays for quantifying the relative binding affinity of DNA sequences for any given TF. These binding assays consisted of multiple *in vitro* experimental methods, such as protein binding microarray (PBM)^41^, HT-SELEX^42^ and SELEX-seq^7^. We previously used these datasets in conjunction with an expanded set of 13 DNA shape features including groove features, inter-base-pair features, and intra-base-pair features (see Methods), demonstrating the effectiveness of DNA shape features in predicting TF-DNA binding specificities using L2-regularized multiple linear regression^27^. We tested DNA shape predicted by Deep DNAshape against the same datasets to determine if improvements could be detected compared to the original pentamer DNAshape model (Fig. 5). Finally, we showed the potential of Deep DNAshape by processing large genomic-level data.

**Fig. 5.**
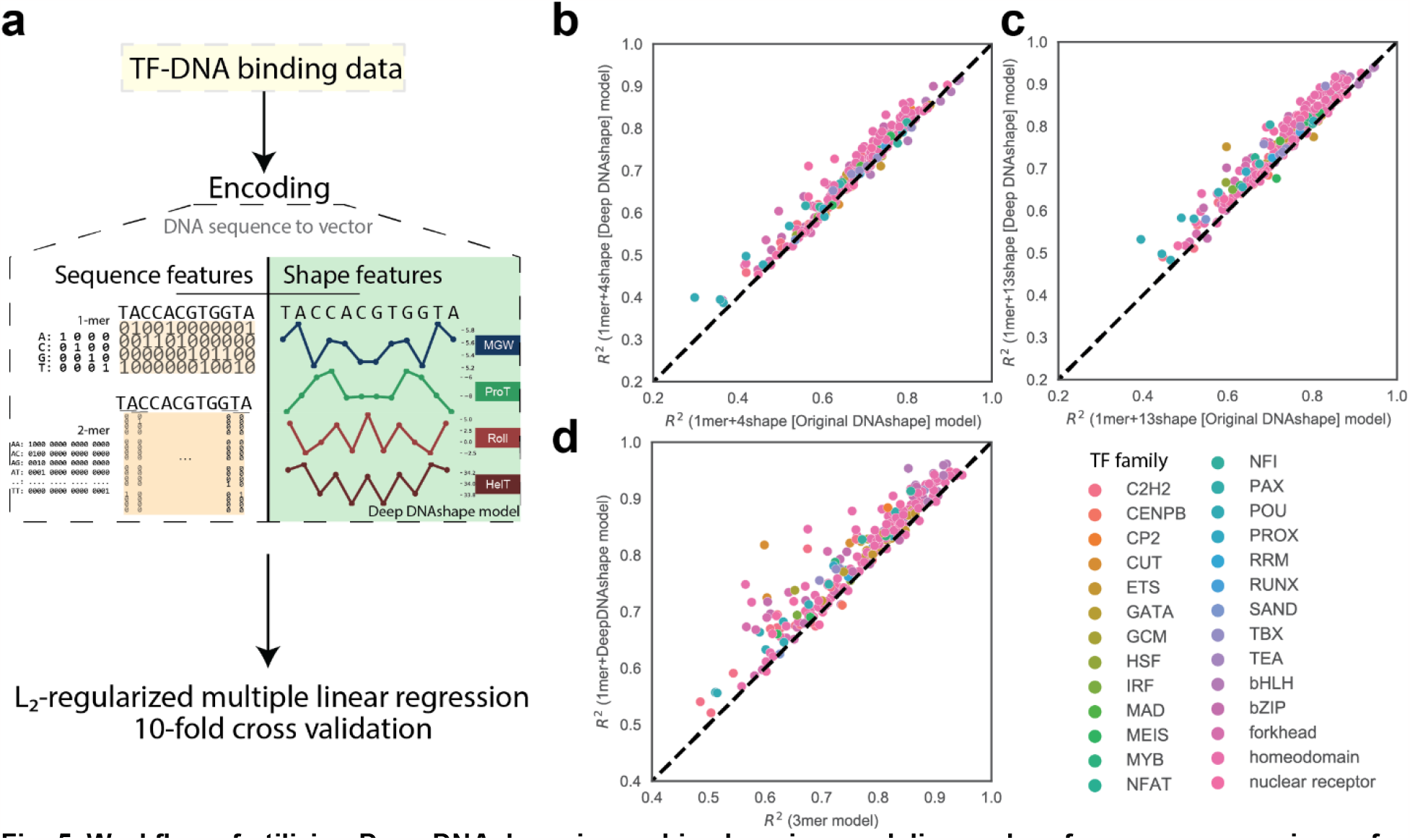
Workflow of utilizing Deep DNAshape in machine learning modeling and performance comparison of binding specificity predictions. **a)** Schematic representation of predicting TF-DNA binding specificities with L2-regularized multiple linear regression model, using encoded sequence features and DNA shape features. **b-d)** *In vitro* TF-DNA binding experimental data used in these models include gcPBM, SELEX-seq and HT-SELEX. **b)** *R*^2^ performance comparison of 1mer+4shape models between original pentamer DNAshape and Deep DNAshape. **c)** *R*^2^ performance comparison of 1mer+13shape models between original pentamer DNAshape and Deep DNAshape. **d)** *R*^2^ performance comparison between 1mer+13shape+FL (the full Deep DNAshape) model and 3mer model without using DNA shape features.

## Results

### Deep DNAshape predicts DNA shape and shape fluctuations considering extended flanking influences without biases

MC simulations were used to generate 3D structural predictions for numerous DNA samples with analyzed DNA shape values for 2,121 different sequences^26^. For each individual DNA shape feature, a training file containing all sequences and their sequence–position-wise DNA shape values was compiled from the simulation data. Despite the varying lengths of these sequences, the Deep DNAshape model was designed to accommodate such variation (Figs. 1a-d, S1). Following hyperparameter searches on training and validation data (Figs. S2, S3), each model was trained on the entire training data and, when used together, was able to predict any DNA shape feature for any length of DNA sequence. The predictions consider only the local base pair information of nearby base pairs (Fig. S1). The models can predict DNA shape features by considering up to 7-bp flanking base pairs, and the number of base pairs of flanking regions considered can be selected for optimal accuracy (Fig. 1e-g). Models were trained with minimal overfitting, while maintaining high accuracy in deeper layers, as evident from the training and validation split samples in the hyperparameter searches (Figs. S2, S3).

We validated our predictions against tetramers compiled from MD simulations^15^ (Table S1) and our previous pentamer query table (Table S2). Additionally, we validated our predictions by calculating and comparing the average inter-base-pair features for 10 di-nucleotides and intra-base-pair features for A-T and C-G base pairs, considering all possible flanking regions predicted by the Deep DNAshape model and its variants using different data sources (Fig. S4, Table S3; also refer to Supplementary Text). This new approach eliminated the use of a query table and enabled us to predict DNA shape features affected by longer flanking regions, compared to using a forcibly generated hexamer or heptamer query table with large numbers of missing values (Figs. 2a-d, S5-S7, Fig. S8 for Deep DNAshape (MD)). Using our new Deep DNAshape method, the inferred DNA shape values based on effects of extended flanking regions (Figs. 2a-d, S5-7) are almost perfectly aligned (Table S2) with statistically computed values. Compared to an interpolated query table, our method corrects the biases from different distributions of *k*-mers and artifacts from molecular simulations or experiments using DNA fragments (Fig. S9).

By performing high-throughput prediction of DNA shape features considering extended flanking regions, Deep DNAshape can be used to investigate the effects of flanking regions on the core DNA structure without requiring MD simulations or XRC experiments. We initially examined four DNA shape features, minor groove width (MGW), propeller twist (ProT), Roll, and helix twist (HelT), at the core of a DNA fragment for all DNA sequence combinations, with the goal of examining the general neighboring effect on different cores (Fig. S10). The static shape of the core can show bimodal or even trimodal distributions, affected by different flanking sequences (Fig. S10).

One DNA fragment worth studying is the A-tract, which contains at least three consecutive ApA, ApT, or TpT bp steps without a flexible TpA bp step, and has been associated with a narrower MGW^1,9^. We permutated all *k*-mers with certain sequences as their cap, such as AAAA-NNNNNNN-AAAA, for 15-mers with A-tract caps. Next, we predicted the MGW using Deep DNAshape compared to counterparts capped by GCGC ends. This approach permitted us to see effects from different flanking cap combinations (Fig. 2e-f). The result indicated that A-tracts increase the flanking MGW on their 5’ end and decrease the MGW on their 3’ end. In other words, the MGW for a short strand of DNA will be upregulated in the 3’ end and downregulated in the 5’ end, if accompanied by two A-tracts at the 5’ and 3’ ends. The A-tracts showed a general trend of decreasing the MGW from 5’ to 3’, consistent with previous individual structural studies^43,44^.

The dynamics of DNA sequences can be difficult to describe. In Deep DNAshape, DNA shape fluctuations can be predicted to represent conformational flexibility for an individual DNA molecule. The training data are based on a compiled shape fluctuation dataset from MC simulations. Different base pairs or bp steps have different intrinsic flexibilities. These fluctuation values are comparable to values that are directly computed from MD simulations (Tables S1)^15^. The values contribute to the global bendability, twistability, and prolong-ability of DNA double helical molecules. The effects of A-tracts on fluctuations can also be visualized (Fig. S11). We observed that flanking A-tracts greatly elevate the MGW fluctuations of the cores (Fig. S11), whereas GCGC ends slightly decrease the MGW fluctuations of the cores.

### Deep DNAshape is superior to pentamer DNAshape in elucidating TF-DNA binding

High-throughput prediction of DNA shape can be applied to data from experimental binding assays on TF-DNA binding (Fig. 3a). The original pentamer DNAshape^26–28,37,45^ method lacked the ability to utilize longer-range flanking regions of sequences. Therefore, this method cannot diversify DNA shape changes in the core binding site affected by extended flanking regions (Fig. 3b) that may still contribute to TF binding specificity. For example, the DNA binding affinity of basic-helix-loop-helix (bHLH) enhancer box (E-box) proteins (Fig. 3c) is greatly affected by the flanking regions^46^. The pentamer DNAshape method cannot access a flanking region-associated change in the DNA shape of the central ‘CG’ di-nucleotide in the most common E-box motif of ‘CACGTG’. In our new model, a ‘shape layer’ capable of utilizing extended flanking regions (Fig. 3b) may provide additional insights, such as into how bHLH proteins bind to their target DNA sequences.

We investigated genomic-context PBM data for three human bHLH TFs, Max, c-Myc, and Mad2. We plotted MGW predictions for the top 25% of aligned binding data (Figs. 3d, S12-13) and compared them to the predictions by the original pentamer DNAshape. Subtler differences can be seen in the central motif cores than was possible with the pentamer-based DNAshape method, even though most of the cores had the same sequence identity ‘CACGTG’ (Fig. 3e-h). When we filtered out the TF-DNA binding dataset to include only sequences with ‘CACGTG’ in the core, we observed a consistent negative correlation between the binding affinities and the Roll and Roll-FL values predicted by Deep DNAshape (Fig. S14), regardless of the *in vitro* experiment types. These correlations are also observable using Deep DNAshape (Expt) (Fig. S15a-c), but not when using Deep DNAshape (MD) (Fig. S15d-f). Such visualizations had been unachievable with the previous pentamer DNAshape method because this method only considers 2-bp flanking regions (which, in the present case, remain constant). These results highlight the effect of the flanks and the need for a method that accounts for more than the nearest- and next-nearest neighbors. Through Deep DNAshape, we can propose a hypothesis for the potential binding mechanism used by the bHLH family to distinguish identical binding cores (Fig. S16), although this hypothesis will require further investigation to confirm.

We also investigated some of the well-studied homeodomain (Hox) proteins (Fig. 4). The Hox proteins Extradenticle (Exd) and Sex combs reduced (Scr) heterodimerize to bind to DNA sequences. The resulting Exd-Scr heterodimer (Fig. 4a-c) prefers a narrower MGW of the two ‘AY’ steps in its motif ‘NGAYNNAY’^7^. Another heterodimer, Exd with Ultrabithorax (Ubx), does not have this preference in the second ‘AY’ step^7^ (Fig. 4d-f). Exd itself has another arginine in the N terminal immediately before the motif that inserts into the DNA minor groove^9^. Compared to the original pentamer-based DNAshape, Deep DNAshape can predict DNA shape features while considering longer flanking regions and in the terminal regions of DNA target sites. Using Deep DNAshape, we revealed a narrower MGW at correct locations according to co-crystal structures using the aligned 16-mer SELEX-seq data (Fig 4b-c, e-f). Furthermore, for Scr, hypotheses can be made that the first ‘AY’ dip in the MGW is required for binding, which is seen across high to low binding sites. The binding affinity is strengthened by a second ‘AY’ dip and by the other dip for Exd binding with its N-terminus. Unlike the pentamer-based DNAshape method, Deep DNAshape predicted the first ‘AY’ dip in the MGW on both A and Y bases. This observation is well-aligned with other Hox protein research on the influence of DNA shape^6^.

### Deep DNAshape exhibits improved prediction accuracy for TF-DNA binding specificity

The mechanisms of TF-DNA readout are complex, leading to substantial research on the use of machine learning to improve the prediction accuracy for TF-DNA binding specificities^27,47,48^. In previous studies, we have successfully improved the performance of multiple linear regression tasks on TF-DNA binding affinity data by incorporating DNA shape features^26,27^ (Fig. 5a). This DNA shape encoding approach reduced the degrees of freedom when using *k*-mer encoding, while maintaining the same level of performance. These models utilized both sequence and DNA shape features to achieve improved performance.

With the introduction of Deep DNAshape, which considers longer-range effects, we can now replace the DNA shape features in these machine learning models with new ones predicted by Deep DNAshape. The DNA shape features predicted by Deep DNAshape outperform the pentamer DNAshape version (Fig. 5b-c). Furthermore, by using the new DNA shape features in combination with fluctuation values, we were able to surpass the performance of the 3-mer model with fewer degrees of freedom (Fig. 5d). Additionally, the updated fluctuation values greatly improved the model compared to the previous SD values^27^ (Fig. S17). From a machine learning perspective, the DNA shape and fluctuation values predicted by Deep DNAshape contain more relevant information. We subsequently compared Deep DNAshape with its variants trained by different underlying data sources, finding that their performances were relatively similar (Fig. S18). Deep DNAshape retains its performance in deeper layers that consider longer flanking regions, while the variants peak at layer 2 or 3 (See Supplementary Text).

### Deep DNAshape reveals a more conserved relationship of DNA shape in transcription start sites (TSSs) between fly species

One key advantage of the DNAshape method is its ability to perform high-throughput predictions that can be easily applied to genomic-level data^49^. To assess the performance of Deep DNAshape on a large dataset, we used a dataset of TSSs from four *Drosophila* species (*D. melanogaster, D. simulans, D. sechellia*, and *D. pseudoobscura*)^50^. Previous research highlighted the importance of structural features in protein binding at TSSs^51^ . Our analysis of DNA shape features at TSSs for these four fly species revealed conserved relationships in DNA shape features among these genomic regions despite sequence differences^49^. Data for this analysis were derived from gene expression experiments^50^, and Deep DNAshape was able to process these data in a matter of minutes on a machine equipped with an NVIDIA A100. Our results showed more evolutionarily conserved relationships in the genomic structural features across these four fly species, as evidenced by the MGW, ProT, Roll, and HelT values (Fig. S19, Table S4).

## Discussion

In this study, we introduce Deep DNAshape, a high-throughput method that accurately predicts 3D DNA structural parameters, or DNA shape, for any DNA sequence. Our previous high-throughput prediction method, DNAshape, successfully predicted DNA shape features, including MGW, inter-base-pair features, and intra-base-pair features^27,37^, and was useful for studying structural TF-DNA binding mechanisms without the need of extensive molecular simulations or structural biology experiments. However, it had several major drawbacks and limitations.

Firstly, DNAshape relied entirely on a pentamer query table to predict DNA shape features at central positions. Thus, to predict the DNA shape feature for the pentamer core, only two bp of flanking regions were considered. The query table could not cover the full two bp of possible flanking combinations for inter-base-pair features, and any sequences in the flanks beyond two bp were not included in predictions. This query table may be sufficient if flanking regions over two bp away do not contribute to DNA shape at the core of the target sequence at first approximation. However, evidence has shown^46^ that such extended flanking regions may affect the binding affinity and DNA shape at the core of the DNA fragment of interest. Such effects are evident in the training curve (Fig. S2-3) of Deep DNAshape.

Secondly, sequence combinations of the flanking regions outside the pentamers were not evenly distributed in the simulation data, which introduced biases towards each pentamer calculation (Fig. S9). For example, if a pentamer is ‘ACGTA’ and the only heptamer in the simulation data is ‘CACGTAG’, assuming the third base pair away from the center is still affecting the DNA shape in the core, the prediction of DNA shape features for ‘ACGTA’ is always biased towards ‘C’ and ‘G’ flanking the pentamer.

Our proposed Deep DNAshape method (Fig. 1) addresses the limitations of previous methods and significantly improves the ability to predict DNA shape. Unlike the original DNAshape method that relied on pentamer query tables and sliding-window algorithms to assign DNA shape values, Deep DNAshape eliminates the use of these tools. All *k*-mers needed to occur at least once in the simulation data to complete the previous query table, but table completion was not sufficient to prevent statistical bias or artifacts. Even with pentamers that occurred more than 250 times^27^ in the simulation data, the data still did not fully cover all hexamers (Figs. S6, S7), let alone heptamers (Figs. 2a-b, S5), octamers, or longer *k*-mers. Different flanking regions have different effects on the DNA shape features in the core of DNA target site, and these influences may be relevant to sequence components and inferable. Deep DNAshape infers the missing values with the learnt influence of flanking regions on the core, while not harming the prediction considering short flanking regions (Table S2).

Because the method learns the effects of flanking regions and predicts DNA shape features in a layer-by-layer manner, it is self-constrained. Each new layer infers the DNA shape features based on the DNA shape features inferred by its previous layer, while considering one more base pair of flanking regions. Outputs of all layers in the training are used in calculating the loss function, enabling the model to predict DNA shape features considering any length of flanking regions, with near-zero overfitting in deeper layers (Figs. S2, S3). The Deep DNAshape method establishes a general framework for predicting response variables at the single-nucleotide level, considering local environments for variable lengths of DNA, and provides valuable improvements upon the widely used DNAshape method.

DNA shape information is valuable in understanding TF-DNA binding events^1,2,27,46,52–54^. One crucial application of the new Deep DNAshape method is to investigate DNA shape preferences in TF-DNA bindings. With the ability to predict accurate DNA shape features considering longer-range flanking regions, our results provide insight into TF-DNA binding interactions that were previously extremely challenging to investigate (Figs. 3, 4). To quantify the extent to which Deep DNAshape provides more information content than our previous pentamer-based DNAshape method, we applied it to the prediction of TF-DNA binding specificities and found a significant improvement compared to pentamer DNAshape (Fig. 5). The Deep DNAshape method still operates in a high-throughput manner, allowing for the investigation of data on a genomic scale. In a brief application, we demonstrated closer evolutionary relationships in DNA shape parameters between four fly species in TSS regions (Fig. S19). Other recent studies^55–60^ incorporating DNA shape features derived from the pentamer model will very likey benefit from using Deep DNAshape. Thus, the Deep DNAshape method unlocks a whole new level of genomic study of DNA shape.

Besides training the Deep DNAshape model using MC simulations as the underlying data, the model is capable of being re-trained using alternative data sources. Our benchmark analysis revealed that the Deep DNAshape variants Expt (trained by experimentally solved structural data) and MD (trained by MD simulation data) provided similar DNA shape predictions and performance metrics across multiple applications. However, the underlying data of these variants were affected by noise/artifacts or low coverage (see Supplementary Text), making MC simulations the current optimal choice for studying effects of longer flanking regions. Future advancements in MD simulations could allow Deep DNAshape to be easily transitioned to using such data as the underlying source.

In addition to predicting DNA shape, Deep DNAshape is also a general framework for processing variable DNA lengths in a layer-by-layer manner, by using expanded neighbor information of DNA sequences and by inferring response variables (e.g., nucleosome positionings or 3D genome interactions) at the single-nucleotide level. Other research that utilizes DNA shape as a feature would benefit from the use of the Deep DNAshape method. To further improve Deep DNAshape, one could focus on generating additional simulation data, optimizing the DNA shape inference equation or network architecture, and expanding the model to include chemically modified base pairs or mismatched base pairs^11^, if such data could be acquired on a large scale.

## Methods

### DNA structural simulations and DNA shape analyses

DNA sequences with variable lengths are initialized and simulated by the Monte-Carlo (MC) method^26^. After filtering out artifacts, the total number of valid DNA simulations is 2,121^26^. DNA shape features are then calculated by Curves 5.3^12^ and minor groove width (MGW) is symmetrized with respect to the base pair^26^. Data are pooled into a training file that includes all DNA sequences and their corresponding DNA shape values. DNA shape fluctuation values are also calculated by Curves 5.3 through analyzing the sampling process from the MC simulations. Fluctuation values represent the variance that occurred during simulations. These values can be viewed as a correlated measure of DNA conformational flexibility of an individual DNA molecule. In this article, the addition of ‘-FL’ to a DNA shape feature indicates the fluctuations (FL) of that specific DNA shape feature. Note that DNA shape can be derived from other data sources (See Supplementary Methods).

### Definitions of DNA shape features

DNA shape features are used to describe the DNA structure numerically, in a base-pair to base-pair manner. The set of DNA shape features used includes six inter-base-pair features, six intra-base-pair features, and two minor groove features. Additional DNA shape features include base-pair axis features and backbone torsion features, which can be analyzed and predicted if supplied to the model. Inter-base-pair features are used to describe translational distances (in Å) and rotational angles (in °) between adjacent base pairs. Specifically, the six inter-base-pair features are ‘Shift’, ‘Slide’, ‘Rise’, ‘Tilt’, ‘Roll’ (Fig. 1g), and ‘Helix twist (HelT)’ (Fig. 1g). The intra-base-pair features describe the translational distances and rotational angles between two bases in a single base pair. The six intra-base-pair features are ‘Shear’, ‘Stretch’, ‘Stagger’, ‘Buckle’, ‘Propeller twist (ProT)’ (Fig. 1g), and ‘Opening’. The minor groove features describe the groove geometry and electrostatic potential in the center of the minor groove. The groove features used in this article are ‘Minor groove width (MGW)’ (Fig. 1g) and ‘Electrostatic Potential (EP)’^28^. For detailed information, refer to^27^.

### Pre- and post-processing of DNA shape values

To ensure robustness in our DNA shape analysis, we utilized a normalization method to account for different ranges of minimum and maximum values of DNA shape features. We compensated for extreme values (e.g., from simulation artifacts) by using the following normalization equation:

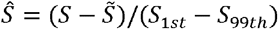

Here, *S* represents the DNA shape feature analyzed from the MC simulation. 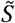 denotes the median of the DNA shape feature values within the dataset, while *S*_1*st*_ and *S*_99*th*_ mark the first percentile and last percentile of the sorted DNA shape feature values, respectively. After normalization, *Ŝ* may still exceed the range of -1 to 1 for extreme values. However, during the training process, the tanh activation function enforces a strict upper and lower bound of *S*_1*st*_ and *S*_99*th*_, ensuring that the output remains within a normalized range of -1 to 1. In postprocessing of the model output, given the normalized DNA shape values predicted by our model, *S* can be directly computed from *Ŝ* using the reverse of the above equation given knowledge of 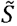, *S*_1*st*_, and *S*_99*th*_.

### Definition of the DNA shape layer

The DNA shape layer is used to treat linear DNA sequences as double-linked nodes or a graph, where each node can be a single nucleotide or di-nucleotides. Each node is connected by its 5’ node and 3’ node through a forward and a backward ‘bond’ (edge). To compute the output features 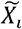 of the shape layer, features are gathered for each node from the previous node (*X*_*i*−1_), current node (*X*_*i*_), and next node (*X*_*i*+1_). 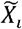 is computed through the following equation,

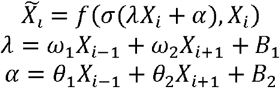

*f* is a trainable gated recurrent unit (GRU) cell with a sigmoid recurrent activation function and tanh activation function. *σ* is ReLu activation followed by batch normalization function. *ω*_1_, *ω*_2_, *θ* _1_, *θ*_2_, *B*_1_, and *B*_2_ are trainable variables to be learnt from the dataset by the model. The self-shape layer is a one-dimensional convolutional layer that transforms the feature number to match later DNA shape layers. This value remains the same for all DNA shape layers.

### Dropout layers and average layers

For each DNA shape layer, a feature vector is generated for every node in the DNA sequence. To prevent overfitting, the feature vector passes through a dropout layer. During prediction, the dropout layer is not used. Finally, the feature vectors are averaged into a single value and post-processed to remove the effects of normalization before making a prediction.

### DNA sequence encoding

To represent the DNA sequences, four characters, ‘A’, ‘C’, ‘G’ and ‘T’ are used, and are one-hot encoded into four arrays as [1,0,0,0], [0,1,0,0], [0,0,1,0] and [0,0,0,1]. For example, a sequence, ‘ACGTGCG’, is represented as

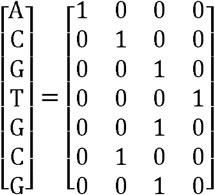

To represent the DNA sequences as di-nucleotides, one-hot encoding for di-nucleotides was used. Each di-nucleotide will be assigned a vector with 16 binary values to represent 16 possible di-nucleotide combinations. The same example ‘ACGTGCG’ will be encoded as

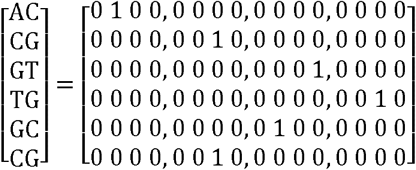

An unknown nucleotide ‘N’ can also be represented this way, where ‘N’ will be a vector by taking the average values upon all possible values. In the implementation of Deep DNAshape, to balance the degree of freedom of the terminal bases, ‘N’ caps are added to both terminals, but they are removed in the final prediction. These ‘N’ caps have a 3’ bond and a 5’ bond connected to themselves.

### Deep DNAshape model design and learning objectives

The model is designed for step-by-step, expandable learning of DNA shape features for any given DNA sequence, with input as one-hot-encoded DNA sequences and output as predicted DNA shape features for each position on the sequence. Inter-base DNA shape features are encoded as di-nucleotides, and sequences are represented as linear double-linked nodes. Mean absolute error (MAE) is used as the loss function, calculated between predicted and MC-simulated DNA shape values from output layers. A postprocessing step is used to recover DNA shape values from normalization. Individual models are trained for each DNA shape feature, with one ‘self’ convolutional layer for input and seven ‘shape layers’ following it.

### Hyperparameter search

Hyperparameters for the Deep DNAshape model include learning rate, optimizer, filter size, and others. These hyperparameters are grid searched for each DNA shape feature and evaluated on a separate training and validation dataset. The best-performing hyperparameters are selected and applied to all DNA shape features. After hyperparameter searches, the model parameters are set as follows: number of shape layers: 7, learning rate: 0.05, number of epochs: 1500, optimizer: stochastic gradient descent (SGD) with momentum 0.95, dropout ratio: 0.5 and filter size: 64.

### Data of transcription factor (TF)-DNA binding assays

We collected and used relative binding affinity data captured in multiple experiments, the same as were used in^27^. The genomic-context PBM data include human TFs c-Myc, Max, and Mad2^61^, where Max as Max-Max homodimer, c-Myc as c-Myc-Max heterodimer and Mad2 as Mad2-Max heterodimer. HT-SELEX data include many TFs in multiple TF families^62^. SELEX-seq data include TFs in the Hox family from *Drosophila*^5,7^.

### L2-regularized multiple linear regression model for TF-DNA binding prediction

The L2-regularized multiple linear regression model is designed to predict relative TF-DNA binding specificity from the data of the TF-DNA binding assays^27^. The model encodes DNA sequences as *k*-mer (*k*=1,2,3) sequence features and any number of DNA shape features. Unless otherwise specified, ‘4shape’ indicates four shape features of MGW, ProT, Roll, and HelT, and ‘13shape’ indicates the MGW plus the 6 inter-base-pair features and 6 intra-base-pair features. The DNA shape features are normalized according to the minimum, maximum, and standard deviation seen in the dataset. Input data are separated into 10 folds. In each fold of the training and test data, another 10-fold cross validation is used in the training data to select lambda values for the L2 term. The model then uses the selected lambda to fit the training data and predict the values for the test data. In the end, 10 folds of predictions are combined to assess the model performance as *R*^2^. The number of sequence features, for sequences with length *n*, is 4*n* for 1-mers, 16 . (*n* − 1) for 2-mers, and 64 . (*n* − 2) for 3-mers. The number of shape features, for sequences with length *n*, is *n* for each base-pair feature and groove feature, and *n* − 1 for each base-pair step feature.

## Supporting information

Supplementary Text, Methods, Tables, Figures, and References

## Author Contributions

J.L., T.C. and R.R. conceived the project. J.L. and T.C. designed the model architecture. J.L. performed all training, evaluations, and analyses. J.L. wrote the manuscript with help from T.C. and R.R. R.R. supervised the project.

## Data availability statement

TF-DNA binding datasets were derived from public resources (see Methods, Data of transcription factor (TF)-DNA binding assays). Raw data for the underlying training datasets (MC, MD and Expt) were sourced from reference^26^ and public databases (see Supplementary Methods for details).

## Code availability statement

All code related to training, prediction, and the pretrained models – as well as an executable package for predicting DNA shape features from any DNA sequence – can be found at https://github.com/JinsenLi/deepDNAshape.

## Acknowledgements

This work was supported by the National Institutes of Health [grant R35GM130376 to R.R.] and the Human Frontier Science Program [grant RGP0021/2018 to R.R.].

## References

1. Rohs, R. et al. Origins of specificity in protein-DNA recognition. Annu. Rev. Biochem. 79, 233–269 (2010).

2. Inukai, S., Kock, K. H. & Bulyk, M. L. Transcription factor–DNA binding: beyond binding site motifs. Curr. Opin. Genet. Dev. 43, 110–119 (2017).

3. Paillard, G., Deremble, C. & Lavery, R. Looking into DNA recognition: zinc finger binding specificity. Nucleic Acids Res. 32, 6673–6682 (2004).

4. Siggers, T. W. & Honig, B. Structure-based prediction of C2H2 zinc-finger binding specificity: sensitivity to docking geometry. Nucleic Acids Res. 35, 1085–1097 (2007).

5. Abe, N. et al. Deconvolving the Recognition of DNA Shape from Sequence. Cell 161, 307–318 (2015).

6. Zeiske, T. et al. Intrinsic DNA Shape Accounts for Affinity Differences between Hox-Cofactor Binding Sites. Cell Rep. 24, 2221–2230 (2018).

7. Slattery, M. et al. Cofactor binding evokes latent differences in DNA binding specificity between Hox proteins. Cell 147, 1270–1282 (2011).

8. Kribelbauer, J. F. et al. Context-Dependent Gene Regulation by Homeodomain Transcription Factor Complexes Revealed by Shape-Readout Deficient Proteins. Mol. Cell 78, 152-167.e11 (2020).

9. Rohs, R. et al. The role of DNA shape in protein-DNA recognition. Nature 461, 1248–1253 (2009).

10. Dantas Machado, A. C. et al. Landscape of DNA binding signatures of myocyte enhancer factor-2B reveals a unique interplay of base and shape readout. Nucleic Acids Res. 48, 8529–8544 (2020).

11. Afek, A. et al. DNA mismatches reveal conformational penalties in protein–DNA recognition. Nature 587, 291–296 (2020).

12. Lavery, R. & Sklenar, H. The Definition of Generalized Helicoidal Parameters and of Axis Curvature for Irregular Nucleic Acids. J. Biomol. Struct. Dyn. 6, 63–91 (1988).

13. Pérez, A., Luque, F. J. & Orozco, M. Frontiers in Molecular Dynamics Simulations of DNA. Accounts Chem. Res. 45, 196–205 (2012).

14. Pérez, A., Lankas, F., Luque, F. J. & Orozco, M. Towards a molecular dynamics consensus view of B-DNA flexibility. Nucleic Acids Res. 36, 2379–2394 (2008).

15. Pasi, M. et al. μABC: a systematic microsecond molecular dynamics study of tetranucleotide sequence effects in B-DNA. Nucleic Acids Res. 42, 12272–12283 (2014).

16. Heddi, B., Oguey, C., Lavelle, C., Foloppe, N. & Hartmann, B. Intrinsic flexibility of B-DNA: the experimental TRX scale. Nucleic Acids Res. 38, 1034–1047 (2010).

17. Marin-Gonzalez, A., Vilhena, J. G., Perez, R. & Moreno-Herrero, F. A molecular view of DNA flexibility. Q. Rev. Biophys. 54, e8 (2021).

18. Haran, T. E. & Mohanty, U. The unique structure of A-tracts and intrinsic DNA bending. Q. Rev. Biophys. 42, 41–81 (2009).

19. Nikolova, E. N., Bascom, G. D., Andricioaei, I. & Al-Hashimi, H. M. Probing Sequence-Specific DNA Flexibility in A-Tracts and Pyrimidine-Purine Steps by Nuclear Magnetic Resonance 13C Relaxation and Molecular Dynamics Simulations. Biochemistry 51, 8654–8664 (2012).

20. Ngo, T. T. M. et al. Effects of cytosine modifications on DNA flexibility and nucleosome mechanical stability. Nat. Commun. 7, 10813 (2016).

21. Li, S., Peng, Y., Landsman, D. & Panchenko, A. R. DNA methylation cues in nucleosome geometry, stability and unwrapping. Nucleic Acids Res. 50, 1864–1874 (2022).

22. Ghoshdastidar, D. & Bansal, M. Flexibility of flanking DNA is a key determinant of transcription factor affinity for the core motif. Biophys. J. (2022) doi:10.1016/j.bpj.2022.08.015.

23. Lavery, R., Moakher, M., Maddocks, J. H., Petkeviciute, D. & Zakrzewska, K. Conformational analysis of nucleic acids revisited: Curves+. Nucleic Acids Res. 37, 5917–5929 (2009).

24. Lavery, R. & Sklenar, H. Defining the Structure of Irregular Nucleic Acids: Conventions and Principles. J. Biomol. Struct. Dyn. 6, 655–667 (1989).

25. Lu, X. J. & Olson, W. K. 3DNA: a software package for the analysis, rebuilding and visualization of three-dimensional nucleic acid structures. Nucleic Acids Res. 31, 5108–5121 (2003).

26. Zhou, T. et al. DNAshape: a method for the high-throughput prediction of DNA structural features on a genomic scale. Nucleic Acids Res. 41, W56–W62 (2013).

27. Li, J. et al. Expanding the repertoire of DNA shape features for genome-scale studies of transcription factor binding. Nucleic Acids Res. 45, 12877–12887 (2017).

28. Chiu, T.P., Rao, S., Mann, R. S., Honig, B. & Rohs, R. Genome-wide prediction of minor-groove electrostatic potential enables biophysical modeling of protein–DNA binding. Nucleic Acids Res. 45, 12565–12576 (2017).

29. Barissi, S., Sala, A., Wieczór, M., Battistini, F. & Orozco, M. DNAffinity: a machine-learning approach to predict DNA binding affinities of transcription factors. Nucleic Acids Res. 50, 9105–9114 (2022).

30. Wang, S. et al. Predicting transcription factor binding sites using DNA shape features based on shared hybrid deep learning architecture. Mol. Ther. - Nucleic Acids 24, 154–163 (2021).

31. Demirci, S., Peters, S. A., Ridder, D. & Dijk, A. D. J. DNA sequence and shape are predictive for meiotic crossovers throughout the plant kingdom. Plant J. 95, 686–699 (2018).

32. Zhang, Q., Shen, Z. & Huang, D.-S. Predicting in-vitro Transcription Factor Binding Sites Using DNA Sequence + Shape. IEEE ACM Transactions Comput. Biology Bioinform. 18, 667–676 (2021).

33. Yang, J. et al. Prediction of regulatory motifs from human Chip-sequencing data using a deep learning framework. Nucleic Acids Res. 47, 7809–7824 (2019).

34. Rohs, R., Sklenar, H. & Shakked, Z. Structural and energetic origins of sequence-specific DNA bending: Monte Carlo simulations of papillomavirus E2-DNA binding sites. Structure 13, 1499–1509 (2005).

35. Balaceanu, A. et al. Modulation of the helical properties of DNA: next-to-nearest neighbour effects and beyond. Nucleic Acids Res. 47, 4418–4430 (2019).

36. Rube, H. T., Rastogi, C., Kribelbauer, J. F. & Bussemaker, H. J. A unified approach for quantifying and interpreting DNA shape readout by transcription factors. Mol. Syst. Biol. 14, e7902 (2018).

37. Chiu, T.P. et al. DNAshapeR: an R/Bioconductor package for DNA shape prediction and feature encoding. Bioinformatics 32, 1211–1213 (2015).

38. Young, R. T., Czapla, L., Wefers, Z. O., Cohen, B. M. & Olson, W. K. Revisiting DNA Sequence-Dependent Deformability in High-Resolution Structures: Effects of Flanking Base Pairs on Dinucleotide Morphology and Global Chain Configuration. Life 12, 759 (2022).

39. Ivani, I. et al. Parmbsc1: a refined force field for DNA simulations. Nat. Methods 13, 55–58 (2016).

40. Chiu, T.P., Li, J., Jiang, Y. & Rohs, R. It is in the flanks: conformational flexibility of transcription factor binding sites. Biophys J. 121, 3765–3767 (2022).

41. Berger, M. F. & Bulyk, M. L. Universal protein-binding microarrays for the comprehensive characterization of the DNA-binding specificities of transcription factors. Nature Protocols 4, 393–411 (2009).

42. Jolma, A. et al. DNA-Binding Specificities of Human Transcription Factors. Cell 152, 327–339 (2013).

43. MacDonald, D. et al. Solution structure of an A-tract DNA bend1 1Edited by I. Tinoco. J. Mol. Biol. 306, 1081–1098 (2001).

44. Stefl, R., Wu, H., Ravindranathan, S., Sklenář, V. & Feigon, J. DNA A-tract bending in three dimensions: Solving the dA4T4 vs. dT4A4 conundrum. Proc. Natl. Acad. Sci. U.S.A. 101, 1177–1182 (2004).

45. Rao, S. et al. Systematic prediction of DNA shape changes due to CpG methylation explains epigenetic effects on protein–DNA binding. Epigenet Chromatin 11, 6 (2018).

46. Gordân, R. et al. Genomic Regions Flanking E-Box Binding Sites Influence DNA Binding Specificity of bHLH Transcription Factors through DNA Shape. Cell Rep. 3, 1093–1104 (2013).

47. Zhou, T. et al. Quantitative modeling of transcription factor binding specificities using DNA shape. Proc. Natl. Acad. Sci. U.S.A. 112, 4654–4659 (2015).

48. Alipanahi, B., Delong, A., Weirauch, M. T. & Frey, B. J. Predicting the sequence specificities of DNA- and RNA-binding proteins by deep learning. Nat. Biotechnol. 33, 831–838 (2015).

49. Chiu, T.-P. et al. GBshape: a genome browser database for DNA shape annotations. Nucleic Acids Res. 43, D103–D109 (2015).

50. Main, B. J., Smith, A. D., Jang, H. & Nuzhdin, S. V. Transcription Start Site Evolution in Drosophila. Mol. Biol. Evol. 30, 1966–1974 (2013).

51. Bansal, M., Kumar, A. & Yella, V. R. Role of DNA sequence based structural features of promoters in transcription initiation and gene expression. Curr. Opin. Struc. Biol. 25, 77–85 (2014).

52. Mathelier, A. et al. DNA Shape Features Improve Transcription Factor Binding Site Predictions In Vivo. Cell Syst. 3, 278-286.e4 (2016).

53. Yang, J. & Ramsey, S. A. A DNA shape-based regulatory score improves position-weight matrix-based recognition of transcription factor binding sites. Bioinformatics 31, 3445–3450 (2015).

54. Slattery, M. et al. Absence of a simple code: how transcription factors read the genome. Trends Biochem. Sci. 39, 381–399 (2014).

55. Liu, Z. & Samee, M. A. H. Structural underpinnings of mutation rate variations in the human genome. Nucleic Acids Res. 51, 7184–7197 (2023).

56. Zhang, Y. et al. A novel convolution attention model for predicting transcription factor binding sites by combination of sequence and shape. Brief. Bioinform. 23, (2021).

57. Ding, P. et al. DeepSTF: predicting transcription factor binding sites by interpretable deep neural networks combining sequence and shape. Brief. Bioinform. 24, (2023).

58. Wang, Z., Xiong, S., Yu, Y., Zhou, J. & Zhang, Y. HAMPLE: deciphering TF-DNA binding mechanism in different cellular environments by characterizing higher-order nucleotide dependency. Bioinformatics 39, btad299 (2023).

59. Bhimsaria, D. et al. Hidden modes of DNA binding by human nuclear receptors. Nat. Commun. 14, 4179 (2023).

60. Khan, S. R., Sakib, S., Rahman, M. S. & Samee, Md. A. H. DeepBend: An interpretable model of DNA bendability. iScience 26, 105945 (2023).

61. Mordelet, F., Horton, J., Hartemink, A. J., Engelhardt, B. E. & Gordân, R. Stability selection for regression-based models of transcription factor-DNA binding specificity. Bioinformatics 29, i117–i125 (2013).

62. Yang, L. et al. Transcription factor family-specific DNA shape readout revealed by quantitative specificity models. Mol. Syst. Biol. 13, 910 (2017).

