## Supplementary Text, Methods, Tables, Figures, and References for "Deep DNAshape: Predicting DNA shape considering extended flanking regions using a deep learning method"

#### Training and evaluating Deep DNAshape on alternative datasets

In the primary manuscript, we discussed how to apply the Deep DNAshape model to DNA shape features analyzed from Monte-Carlo (MC) simulations. Based on our observation of Deep DNAshape performance on MC data, we concluded that the model architecture should extend to any data source. Therefore, we compiled an alternative dataset comprising DNA shape features obtained from experimentally solved structures, and a dataset with DNA shape features derived from a limited number of MD simulations, to benchmark Deep DNAshape. We constructed the experimental dataset using a lightly curated list of PDB entries[1] that includes protein-bound DNA structures and irregular DNA structures, among others. We believe our model architecture can efficiently counter any existing biases and artifacts in this dataset. After processing (refer to Supplementary Methods), DNA shape features were normalized and used to train the Deep DNAshape model, resulting in the Deep DNAshape (Expt) variant. Concurrently, we used DNA shape data from the Parmbsc1 database[2], comprising only B-DNA duplexes, to train another Deep DNAshape variant, Deep DNAshape (MD), despite the limited availability of MD data.

We compared DNA shape predicted by our MC-based Deep DNAshape, Deep DNAshape (Expt), and Deep DNAshape (MD) models in several ways, assuming that the trained Deep DNAshape model can infer neighboring effects as accurately as possible. As the raw underlying data differ in sequence components, they could only be compared using constructed query tables, a method previously validated[3].

Initially, we directly compared the ability of the models to predict *k*-mers (MGW and intra-base-pair feature predictions, on pentamers; inter-base-pair feature predictions on hexamers). We calculated correlations of the core DNA shape across all *k*-mers in the same order between models (Table S3). We also compiled average inter-base-pair shape features across these models for the 10 unique dinucleotides (Fig. S4). The Deep DNAshape (Expt) variant aligns almost identically with the underlying experimental PDB data. Deep DNAshape (MD) generally matches the experimental data, barring a slight overestimation of the feature Slide. Although Deep DNAshape underestimates Roll and Slide values compared to experimental data, the Roll values are better correlated with Deep DNAshape (Expt) compared to Deep DNAshape (MD) (Table S3).

Next, we examined whether features predicted by the variants enhance the performance of the shape-augmented TF-DNA binding prediction models (MLR models). Estimating longer-range neighboring effects is quite challenging with noisy or low-coverage data, even with Deep DNAshape's comprehensive control of over overfitting. We compared performances of 1mer+4shape and 1mer+13shape MLR models (Fig. S22). Up to shape layer 4, the performances of Deep DNAshape (Expt), Deep DNAshape (MD), and Deep DNAshape (MC) are statistically indistinguishable. However, when the layer number further increases, the performance of Deep DNAshape remains constant, whereas the performances of the other two variants exhibit changes (MD in 1mer+4shape model and Expt in both models). As per the design of the Deep DNAshape architecture, early layers do not encompass all longer-range neighboring effects, meaning we should focus on layers 3, 4, and 5. On layer 5, Deep DNAshape outperforms variants trained with other data sources. MD simulations also provide the capacity to learn DNA shape fluctuations. However, the performance of Deep DNAshape (MD) for the 1mer+4shape+FL model suffers dramatically in deeper layers (Fig. S22), likely due to the low coverage of the underlying MD data.

#### Discussion

Successful training of Deep DNAshape on structural data acquired from experimentally solved structures and MD simulations confirms the generality and robustness of the Deep DNAshape model architecture. However, experimentally solved structures contain long DNA sequences bound by proteins only in certain circumstances, possibly deforming the DNA structures to a certain extend. An example is the nucleosome structure[4] of which many different copies exist in the PDB. This causes Deep DNAshape (Expt) models to learn incorrect neighboring effects, leading to worse performance in the deeper layers. Furthermore, available data from MD simulations is currently still limited with tremendous missing values (Fig. S8), causing Deep DNAshape (MD) to learn incomplete neighboring effects from these missing values. Use of MC simulations as our underlying shape source represents at this point a favorable balance of data quality and quantity. Transitioning to MD simulations for training Deep DNAshape will likely become an alternative in the future, provided a sufficiently larger number of MD simulations could be procured.

### Supplementary Methods

#### Acquisition, analysis of experimentally solved structures, and pre-processing for Deep DNAshape (Expt)

Biological assemblies of PDB IDs mentioned in reference[1] were downloaded from the PDB. Duplex DNA helical sections were extracted from all PDB files using X3DNA[5]. DNA shape features were calculated by Curves 5.3, where applicable, resulting in a total of 3,204 PDB IDs (4,034 for biological assemblies). Some bases for which DNA shape could not be computed resulted in NaN values. Numerical shape features in regions containing the abnormal B-DNA helix were set to NaN. Criteria were based on the original DNAshape paper[6], in which regions were disregarded if one of the following conditions existed within 3bp: (i) MGW > 8.5 Å or < 1.5 Å; (ii) HelT > 45°; and (iii) |Roll| > 20°. Chemically modified bases were converted to the original regular base in the final file. Any other irregular bases present in the dataset were set to unknown (N). Shape features of both strands were extracted. Calculated DNA shape features, including features of the reverse complements, were merged. If more than one copy of DNA exists, the values were averaged and NaN values were ignored. This protocol was adopted from[6].

Structures may have very short DNA sequences resulting in more incomputable groove or inter-base-pair features than intra-base-pair features. After preprocessing these structures and merging repeated sequences, we collected 2,129 sequences of available MGW values (about 15.47 bp per sequence), 2,269 sequences of available inter-base-pair shape feature values (about 15.02 bp per sequence) and 2367 sequences of available intra-base-pair shape features (about 14.89 bp per sequence). For comparison purposes, average inter-base-pair shape features are calculated from this training data for each of the 10 unique dinucleotides.

Compiled shape training data are then pre-processed using normalization (see Main Methods). However, due to volatility of this dataset's nature, we adjusted the percentile from 1 to 5:

$$\hat{S} = (S - \tilde{S})/(S_{5th} - S_{95th})$$

Here, $S$ represents the DNA shape feature analyzed from the experimentally solved structural dataset. $\tilde{S}$ denotes the median of the DNA shape feature values within the dataset, while $S_{5th}$ and $S_{95th}$ mark the 5$^{th}$ percentile and 95$^{th}$ percentile of the sorted DNA shape feature values, respectively.

#### Acquisition of MD simulation DNA shape data

DNA shape data files, labeled with tags of “DNA” (DNA only) and “duplex”, were downloaded from <https://mmb.irbbarcelona.org/ParmBSC1/>. Files were merged and aggregated with the addition of reverse complements. We collected 137 sequences for MGW (about 15.89 bp per sequence), 139 sequences containing inter-base-pair features (about 15.92 bp per sequence) and 138 sequences containing intra-base-pair features (about 15.92 bp per sequence).

### Supplementary Tables and Figures

**Table S1.** Correlation of inter-base-pair features predicted by Deep DNAshape and acquired from MD simulations, for all 136 tetramers[7]. Tetramer features from Deep DNAshape are calculated by predicting and averaging from all 8-mers.

| Inter-base-pair features | Spearman's rank correlation of static shape | Spearman's rank correlation of shape fluctuation |
|---|---|---|
| Shift | 0.23 | 0.22 |
| Slide | 0.51 | 0.53 |
| Rise | 0.41 | 0.78 |
| Tilt | 0.59 | 0.85 |
| Roll | 0.79 | 0.32 |
| HelT | 0.44 | 0.41 |

**Table S2.** Pearson's correlations of DNA shape features predicted by Deep DNAshape and reconstructed query table from MC simulations. Reconstructed hexamer and 7-mer query tables contain extensive missing values. There are 10 possible di-nucleotides. We show NpN in this table for easier representation. However, the generated query table for hexamers does not account for all di-nucleotides; hence, the lower correlation. The pentamer query table came with the DNAshape method. "Core" means the middle base pair or base-pair step from the *k*-mer query table.

| Shape feature | Against tetramer | Against pentamer | | Against hexamer | Against heptamer | |
|---|---|---|---|---|---|---|
| | Core: NpN | Core: A/T | Core: C/G | Core: NpN | Core: A/T | Core: C/G |
| MGW | N/A | 0.98 | 0.98 | N/A | 0.96 | 0.96 |
| Shift | 0.99 | N/A | | 0.82 | N/A | |
| Slide | 0.98 | | | 0.87 | | |
| Rise | 0.99 | | | 0.97 | | |
| Tilt | 0.99 | | | 0.78 | | |
| Roll | 0.98 | | | 0.68 | | |
| HelT | 1.00 | | | 0.91 | | |
| Shear | N/A | 0.97 | 0.96 | N/A | 0.94 | 0.92 |
| Stretch | | 0.78 | 0.94 | | 0.77 | 0.87 |
| Stagger | | 0.98 | 0.99 | | 0.97 | 0.98 |
| Buckle | | 0.99 | 0.99 | | 0.98 | 0.99 |
| ProT | | 0.99 | 1.00 | | 0.97 | 0.98 |
| Opening | | 0.93 | 0.97 | | 0.87 | 0.92 |

**Table S3**. Pearson's correlations of DNA shape features predicted by Deep DNAshape and its variants. All unique pentamers (512 sequences) and hexamers (2080 sequences) are used to predict DNA shape. Correlations are calculated by the concatenated predictions as a vector given the same order of sequences.

[See Supplementary_table_S3.xlsx]

**Table S4.** Pearson's correlations of average DNA shape predicted by Deep DNAshape or pentamer DNAshape for four different *Drosophila* species.

MGW from Deep DNAshape

| Species | *D. melanogaster* | *D. simulans* | *D. sechellia* | *D. pseudoobscura* |
|---|---|---|---|---|
| *D. melanogaster* | 1.00 | 0.95 | 0.96 | 0.93 |
| *D. simulans* | | 1.00 | 0.99 | 0.97 |
| *D. sechellia* | | | 1.00 | 0.97 |
| *D. pseudoobscura* | | | | 1.00 |

MGW from original pentamer DNAshape

| Species | *D. melanogaster* | *D. simulans* | *D. sechellia* | *D. pseudoobscura* |
|---|---|---|---|---|
| *D. melanogaster* | 1.00 | 0.93 | 0.91 | 0.91 |
| *D. simulans* | | 1.00 | 0.97 | 0.95 |
| *D. sechellia* | | | 1.00 | 0.95 |
| *D. pseudoobscura* | | | | 1.00 |

ProT from Deep DNAshape

| Species | *D. melanogaster* | *D. simulans* | *D. sechellia* | *D. pseudoobscura* |
|---|---|---|---|---|
| *D. melanogaster* | 1.00 | 0.94 | 0.95 | 0.91 |
| *D. simulans* | | 1.00 | 0.98 | 0.95 |
| *D. sechellia* | | | 1.00 | 0.95 |
| *D. pseudoobscura* | | | | 1.00 |

ProT from original pentamer DNAshape

| Species | *D. melanogaster* | *D. simulans* | *D. sechellia* | *D. pseudoobscura* |
|---|---|---|---|---|
| *D. melanogaster* | 1.00 | 0.93 | 0.90 | 0.89 |
| *D. simulans* | | 1.00 | 0.96 | 0.94 |
| *D. sechellia* | | | 1.00 | 0.93 |
| *D. pseudoobscura* | | | | 1.00 |

Roll from Deep DNAshape

| Species | *D. melanogaster* | *D. simulans* | *D. sechellia* | *D. pseudoobscura* |
|---|---|---|---|---|
| *D. melanogaster* | 1.00 | 0.97 | 0.98 | 0.97 |
| *D. simulans* | | 1.00 | 0.99 | 0.99 |
| *D. sechellia* | | | 1.00 | 0.98 |
| *D. pseudoobscura* | | | | 1.00 |

Roll from original pentamer DNAshape

| Species | *D. melanogaster* | *D. simulans* | *D. sechellia* | *D. pseudoobscura* |
|---|---|---|---|---|
| *D. melanogaster* | 1.00 | 0.94 | 0.95 | 0.94 |
| *D. simulans* | | 1.00 | 0.97 | 0.97 |
| *D. sechellia* | | | 1.00 | 0.96 |
| *D. pseudoobscura* | | | | 1.00 |

HelT from Deep DNAshape

| Species | *D. melanogaster* | *D. simulans* | *D. sechellia* | *D. pseudoobscura* |
|---|---|---|---|---|
| *D. melanogaster* | 1.00 | 0.94 | 0.96 | 0.95 |
| *D. simulans* | | 1.00 | 0.98 | 0.96 |
| *D. sechellia* | | | 1.00 | 0.96 |
| *D. pseudoobscura* | | | | 1.00 |

HelT from original pentamer DNAshape

| Species | *D. melanogaster* | *D. simulans* | *D. sechellia* | *D. pseudoobscura* |
|---|---|---|---|---|
| *D. melanogaster* | 1.00 | 0.93 | 0.91 | 0.93 |
| *D. simulans* | | 1.00 | 0.95 | 0.94 |
| *D. sechellia* | | | 1.00 | 0.94 |
| *D. pseudoobscura* | | | | 1.00 |

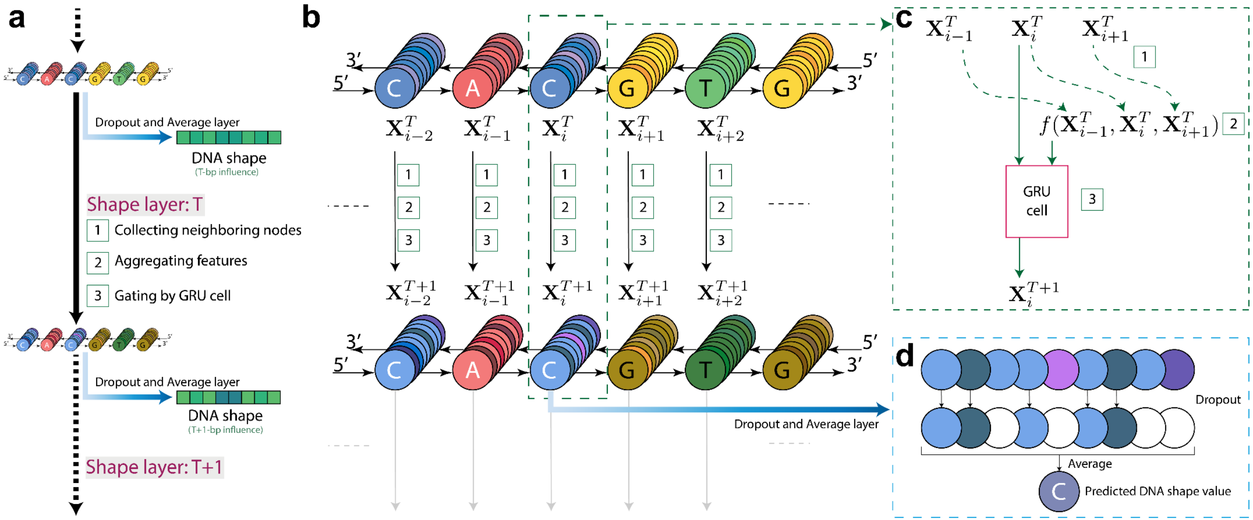


**Fig S1. Deep DNAshape architecture details.**

a) Overview of shape layer. Features (stored in nodes) are used as input to the shape layer. Before the shape layer, nodes are first transformed from one-hot encoding of mono- or di-nucleotides through one self-convolution layer into features. Nodes are interconnected through edges in the data structure. New features (stored in nodes, considering the two neighboring nodes) are the output of the shape layer.

b) Detailed parallel computation schema for shape layer.

c) Calculation in shape layer. Features from neighboring nodes are collected, aggregated by trainable equation (see Main Methods), and gated by a trainable GRU cell to generate the new feature.

d) Dropout and average layer for DNA shape output.

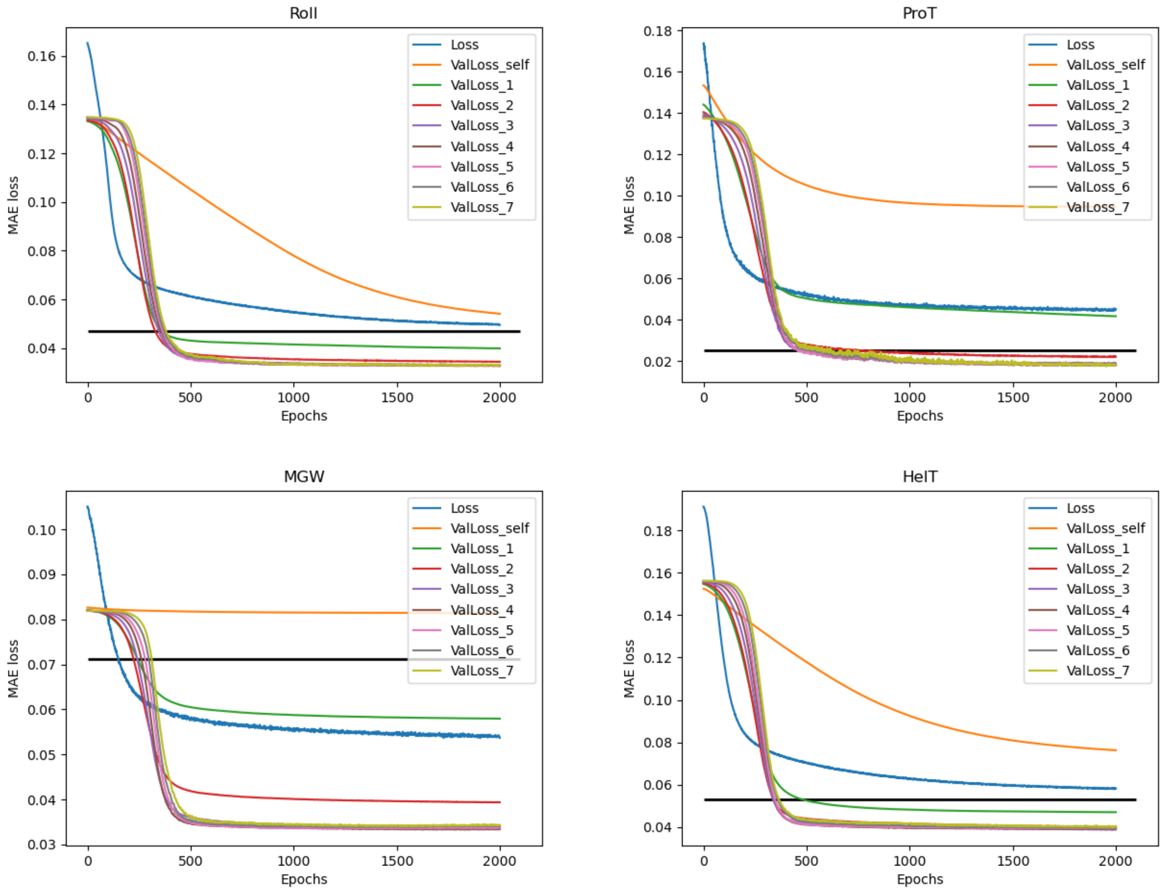


**Fig S2. Training curves for four DNA shape features (Roll, ProT, MGW, and HelT), for a training and validation split (80/20 split).** “Loss” represents training loss of mean absolute error (MAE). “ValLoss” represents the MAE loss on validation set. “self” and “1” to “7” mean different layers outputted from the model. Black horizontal line is a reference loss, as if the same validation data were to be predicted by the original pentamer DNAshape method [3].

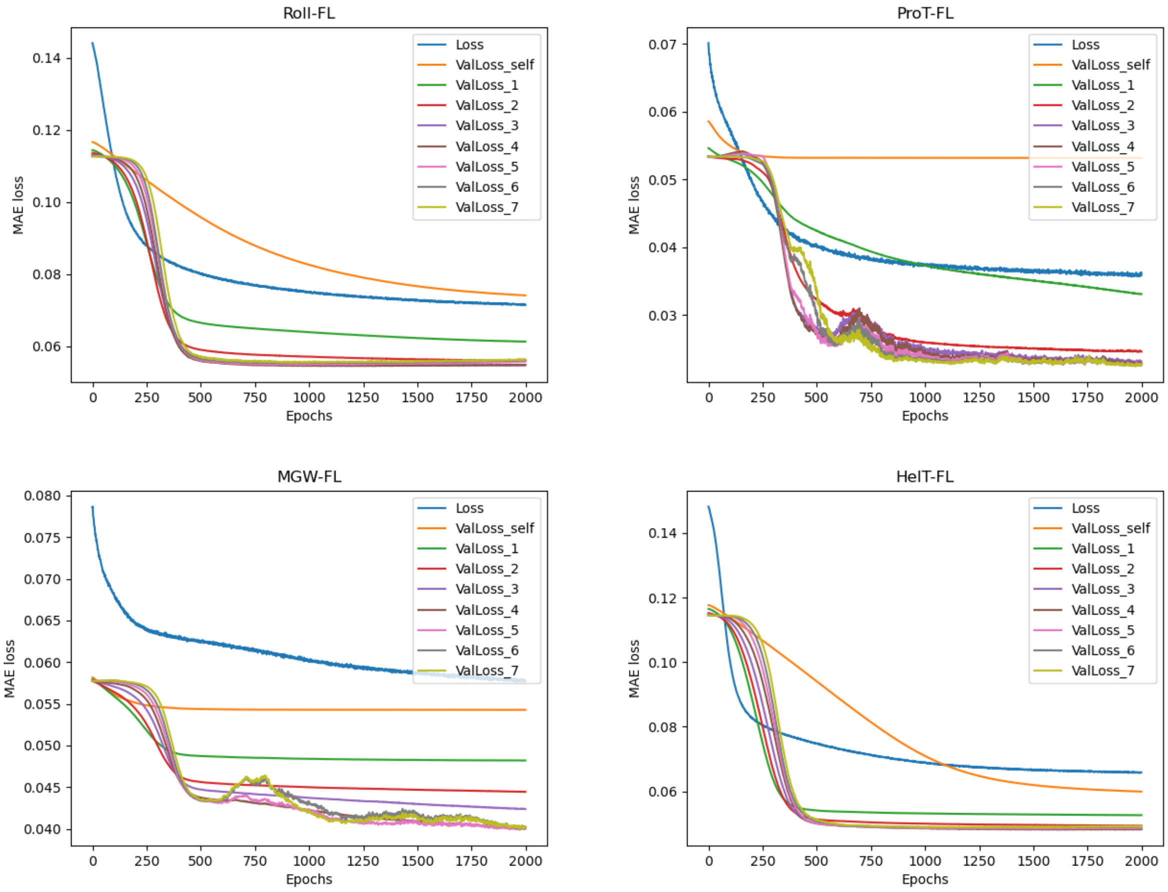


**Fig S3. Training curves for four DNA shape (Roll, ProT, MGW, and HelT) fluctuation values, for a training and validation split (80/20 split).** “Loss” represents the training loss of MAE. “ValLoss” represents the MAE loss on validation set. “self” and “1” to “7” mean different layers outputted from the model. Reference from the original pentamer DNAshape method does not exist.

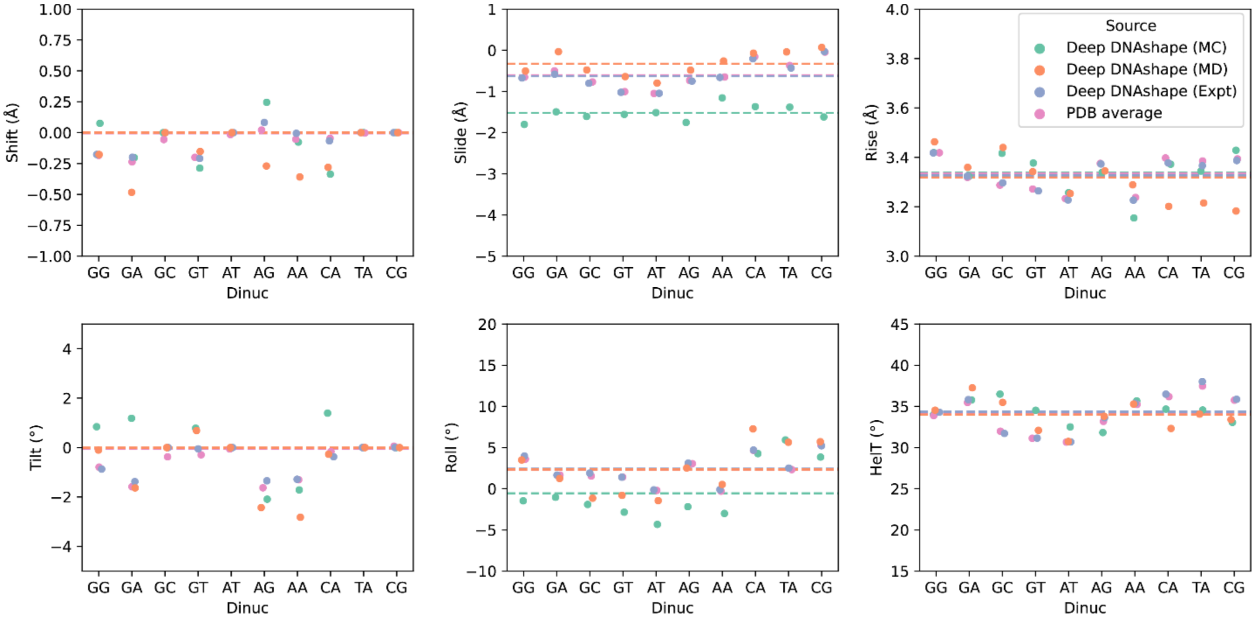


**Fig S4. Averaged inter-base-pair shape features for the 10 dinucleotides.**

Shown are DNA shape features of all hexamers (2,080 sequences) predicted by the Deep DNAshape model and its variants, layer 2. PDB averages are calculated from filtered PDB entries (see Supplementary Methods). Horizontal lines are averaged values among the 10 dinucleotides for each model. Slide and Roll values are underestimated by Deep DNAshape (MC) compared to PDB average. Slide values are overestimated by Deep DNAshape (MD). Values are jittered horizontally to reveal minor difference. Correlations of the 2,080 hexamers can be found in Table S3.

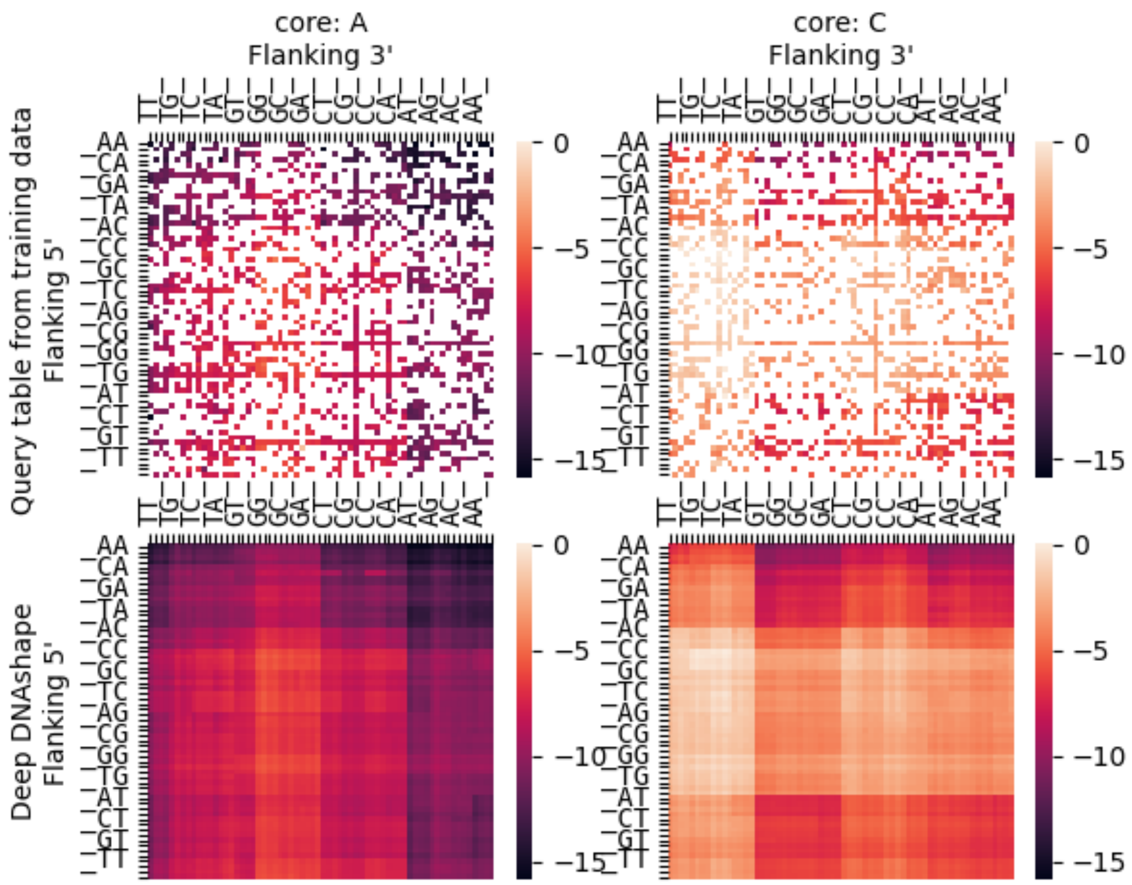


**Fig S5. Heatmaps showing ProT values at central position for all possible 7-mers.** Upper two subpanels show ProT values as if we had constructed a 7-mer query table from available MC simulations directly. Lower two subpanels show ProT values predicted by Deep DNAshape method (layer 3) for all possible 7-mers. Left two subpanels show all sequence with ‘A’ base in the center. Right two subpanels show all sequences with ‘C’ base in the center. “_” represents A, C, G, and T in sequential order. For example, top left grid represents ProT value for AAAATTT in the central position.

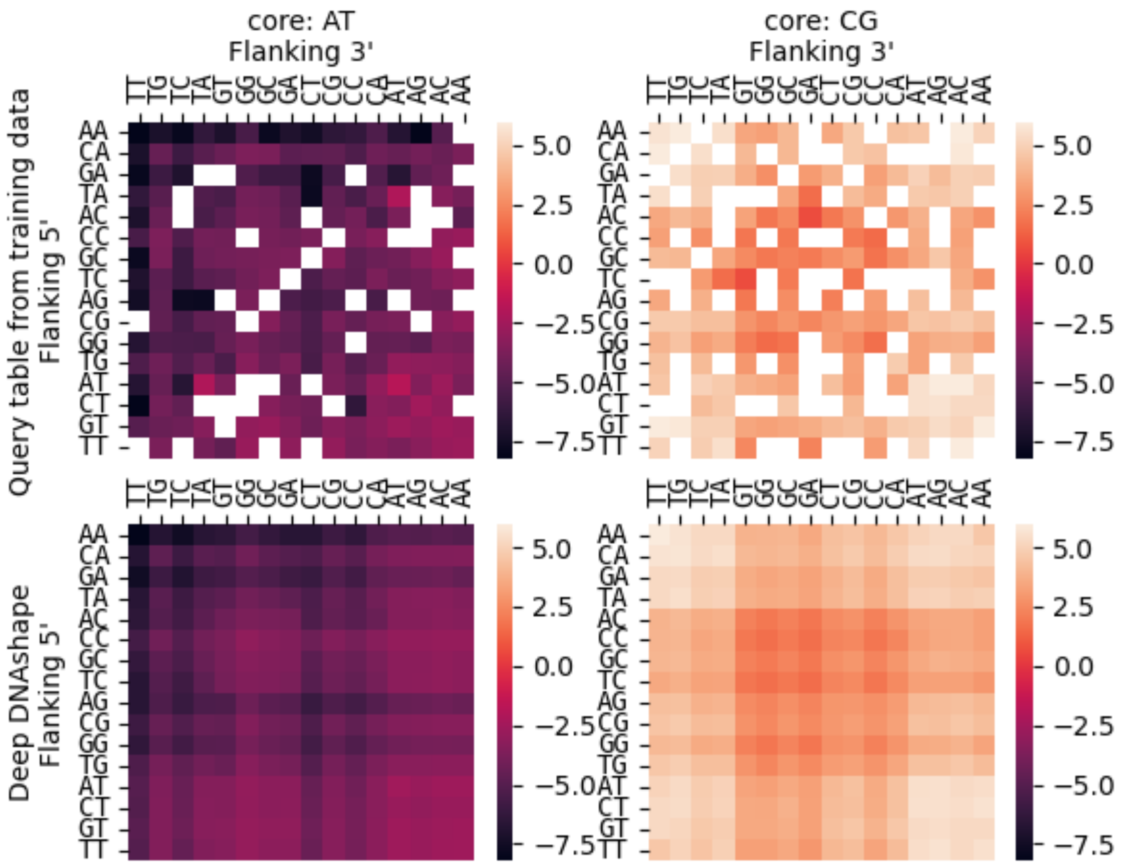


**Fig S6. Heatmaps showing Roll values at central position for all possible 6-mers.** Upper two subpanels show Roll values as if we had constructed a 6-mer query table from available MC simulations directly. Lower two subpanels show Roll values predicted by Deep DNAshape method (layer 3) for all possible 6-mers. Left two subpanels show all sequences with 'AT' dinucleotides in the center. Right two subpanels show all sequences with 'CG' dinucleotides in the center. For example, top left grid represents Roll value for AAATTT in the central position.

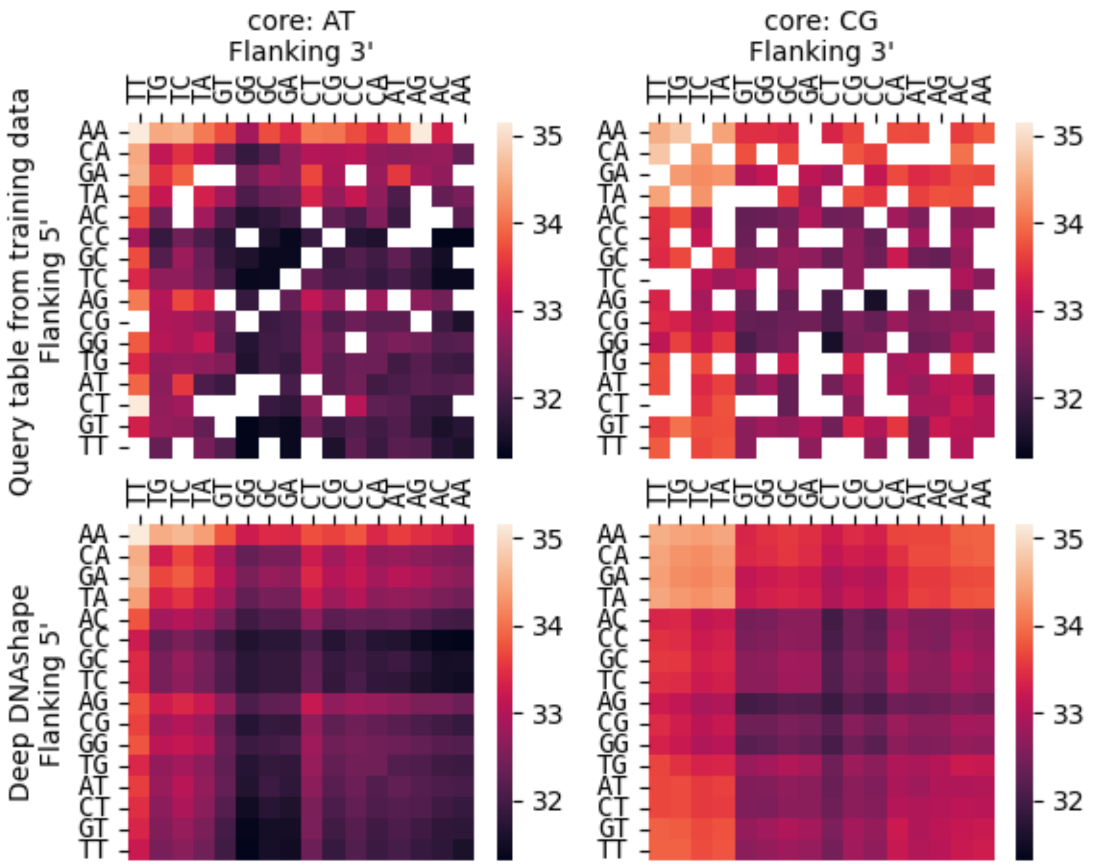


**Fig S7. Heatmaps showing HelT values of middle position for all possible 6-mers.** Upper two subpanels show HelT values as if we had constructed a 6-mer query table from available MC simulations directly. Lower two subpanels show HelT values predicted by Deep DNAshape method (layer 2) for all possible 6-mers. Left two subpanels show all sequences with 'AT' dinucleotides in middle. Right two subpanels show all sequences with 'CG' dinucleotides in middle. For example, top left grid represents HelT value for AAATTT in the central position.

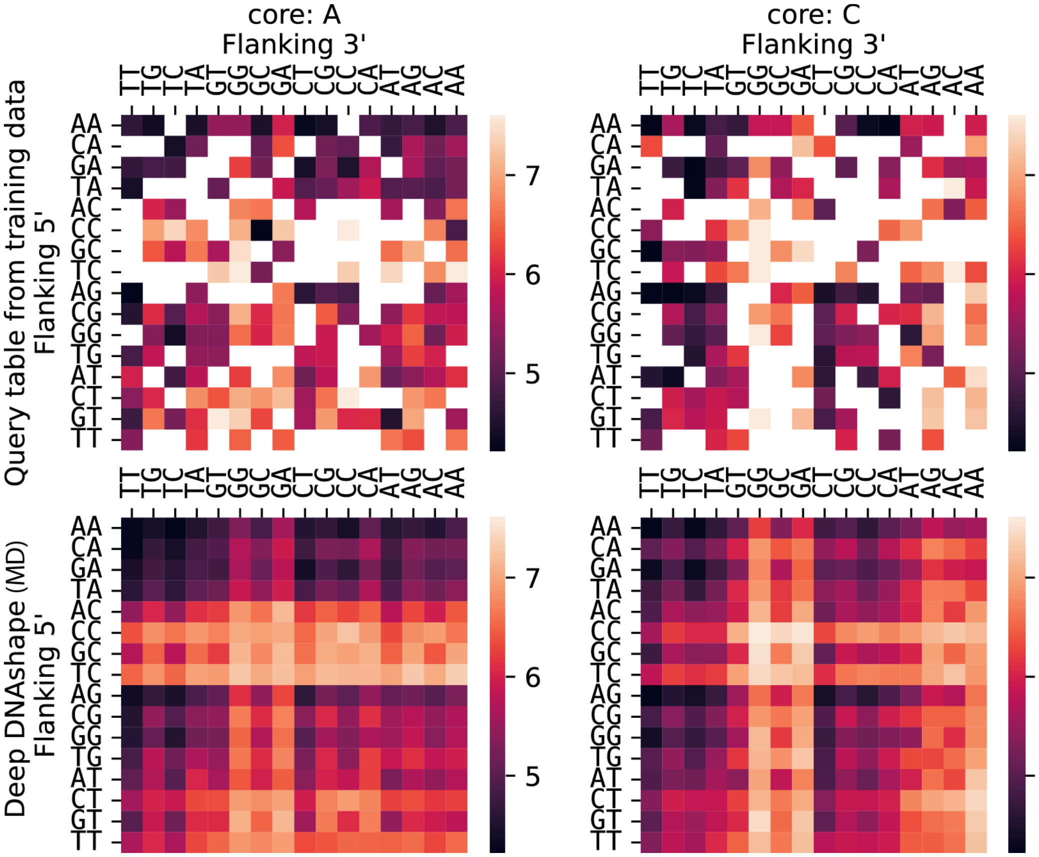


**Fig S8. MD version of heatmaps showing MGW values at central position for all possible 5-mers.** Upper two subpanels show MGW values as if we had constructed a 5-mer query table from available MD simulations directly. Lower two subpanels show MGW values predicted by Deep DNAshape (MD) method (layer 2) for all possible 5-mers. Left two subpanels show all sequences with ‘A’ base in the center. Right two subpanels show all sequences with ‘C’ base in the center.

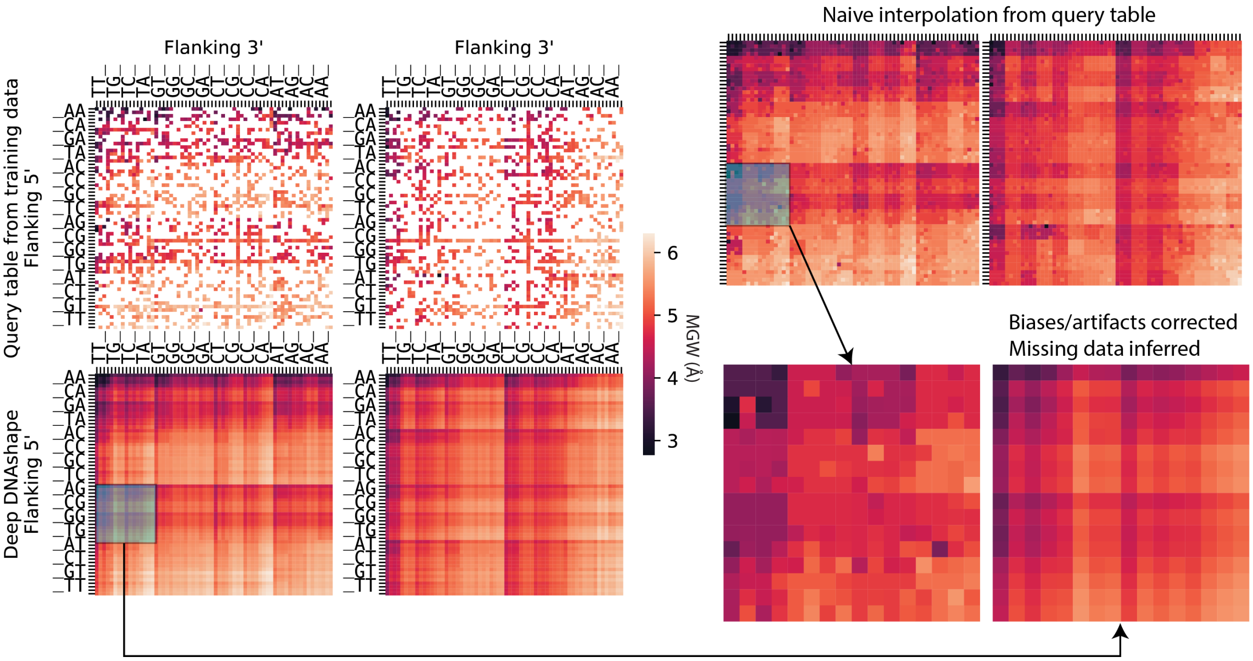


**Fig S9. Comparison between naïve interpolation and Deep DNAshape for generating heptamer query table from training data.** Patch showing MGW for all possible NNGATNN (N can be any of A,C,G or T) sequences is selected for comparison in right bottom panels. Deep DNAshape provides detailed, unbiased information on effects of flanking regions compared to interpolated version.

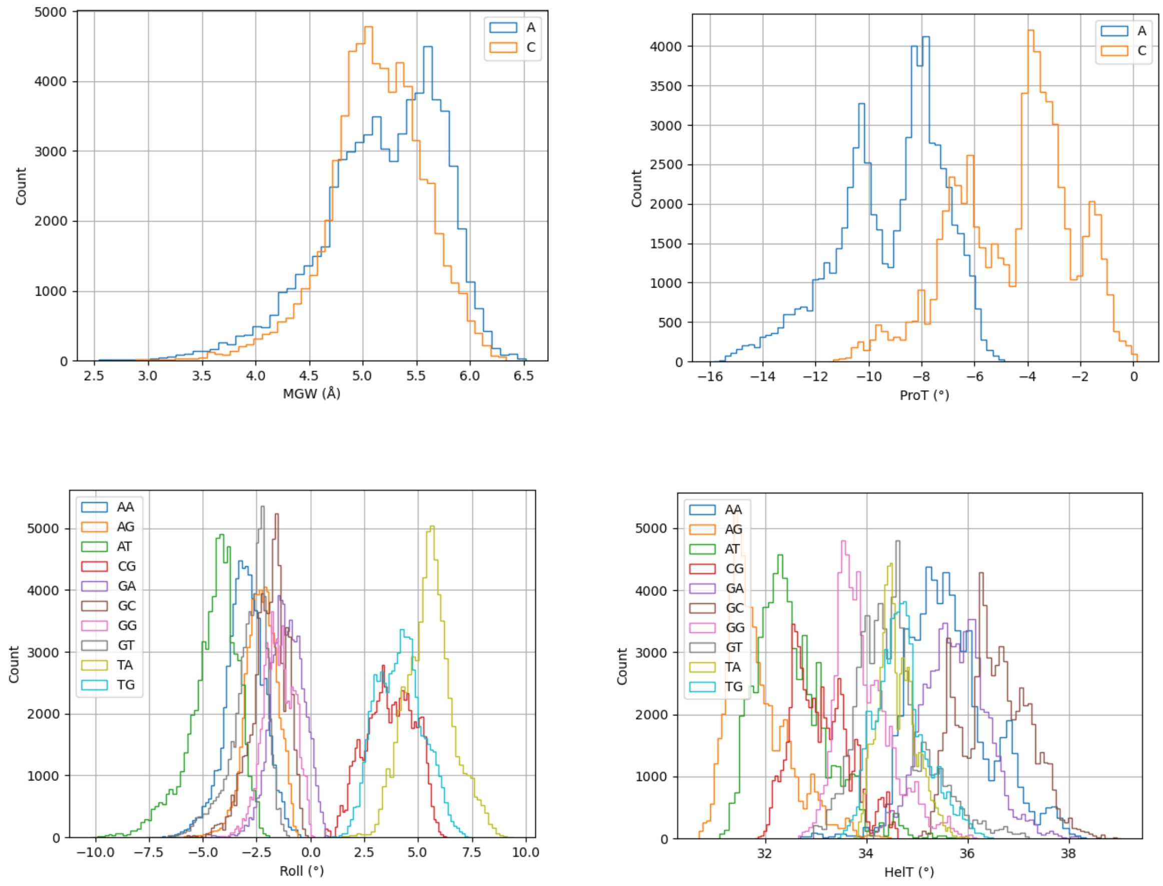


**Fig S10. Predicted DNA shape features for cores (shown in legend), affected by all possible 4-bp of flanking regions.** Features are predicted by Deep DNAshape method (layer 4). MGW is a groove feature assigned to mononucleotides, and ProT is an intra-base-pair feature. Roll and HelT are inter-base-pair features assigned to di-nucleotides. Histograms are generated from all possible 9-mers for MGW and ProT, 10-mers for Roll and HelT.

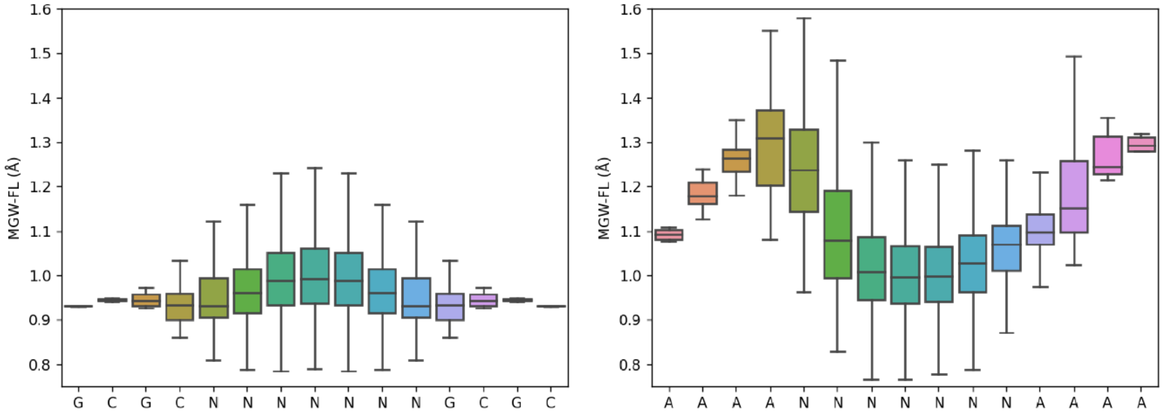


**Fig S11. MGW-FL (minor groove width fluctuations) predicted by Deep DNAshape (layer 4) for all possible random sequences with fixed 5' and 3' caps.** Left panel is capped by 'GCGC'. Right panel is capped by 'AAAA'. Center line indicates the median. Box limits are $75^{th}$ and $25^{th}$ percentiles. The whiskers extend 1.5 times the IQR from the top and bottom of the box. Outliers are removed in boxplots.

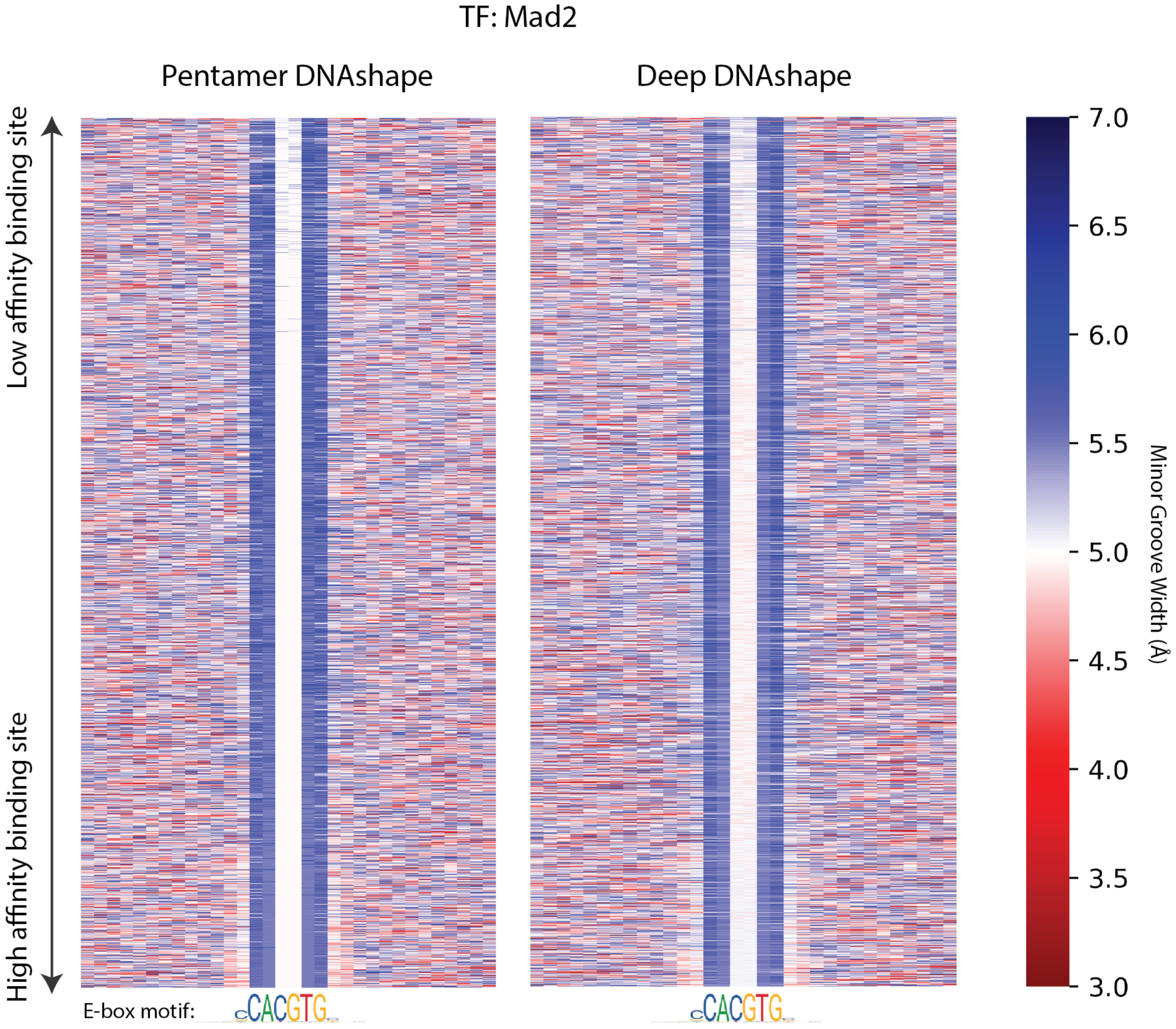


**Fig S12. MGW predicted by original pentamer DNAshape, or by Deep DNAshape (layer 4), for protein Mad2 (Mad2-Max heterodimer) and DNA binding data, in order of relative binding affinity.** Color represents DNA shape values. Data are aligned with core binding site. Only top 25% of binding data are used.

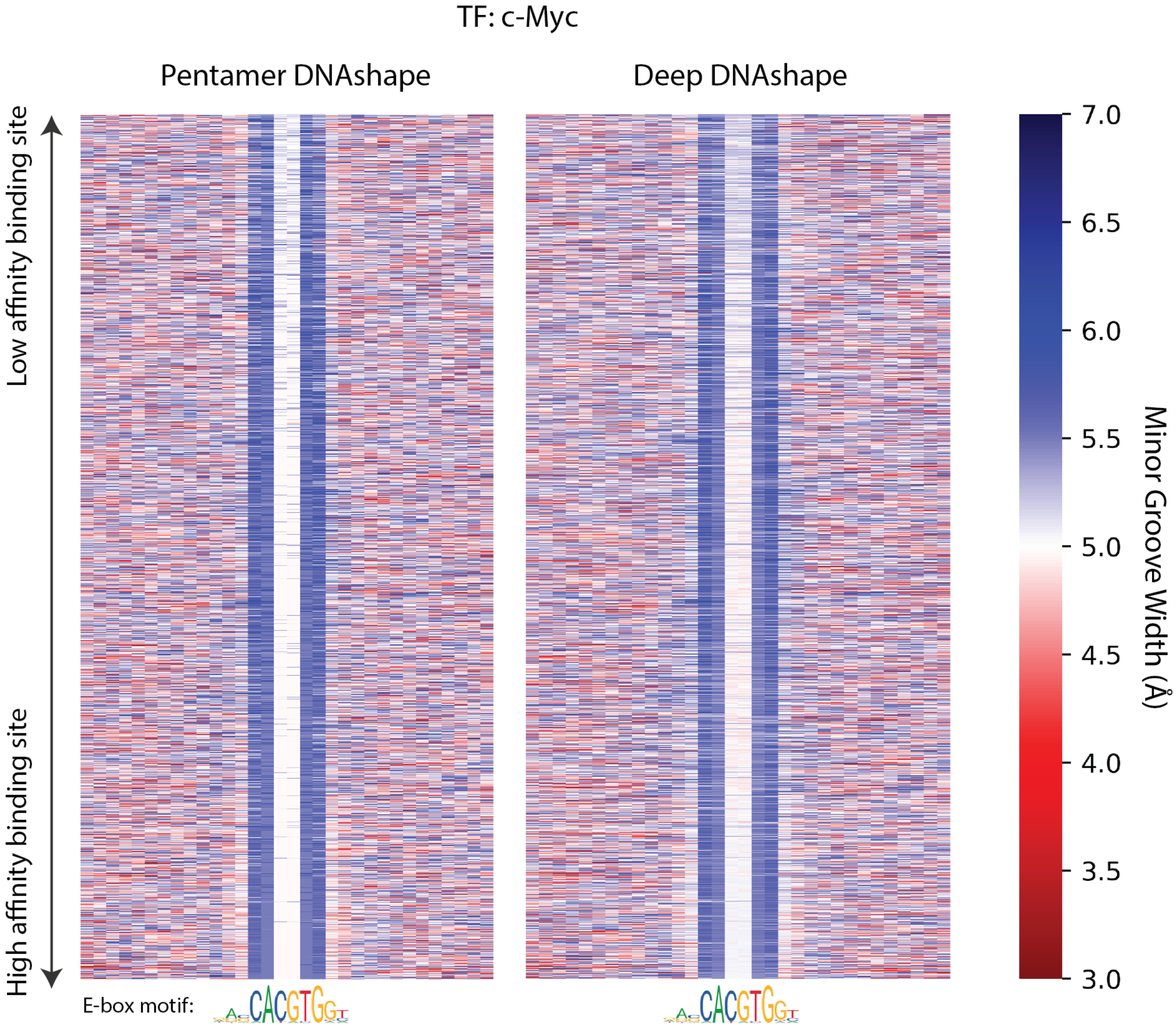


**Fig S13. MGW values predicted by original pentamer DNAshape or by Deep DNAshape (layer 4), for protein c-Myc (c-Myc-Max heterodimer) and DNA binding data, in order of relative binding affinity.** Color represents DNA shape values. Data are aligned with core binding site. Only top 25% of binding data are used.

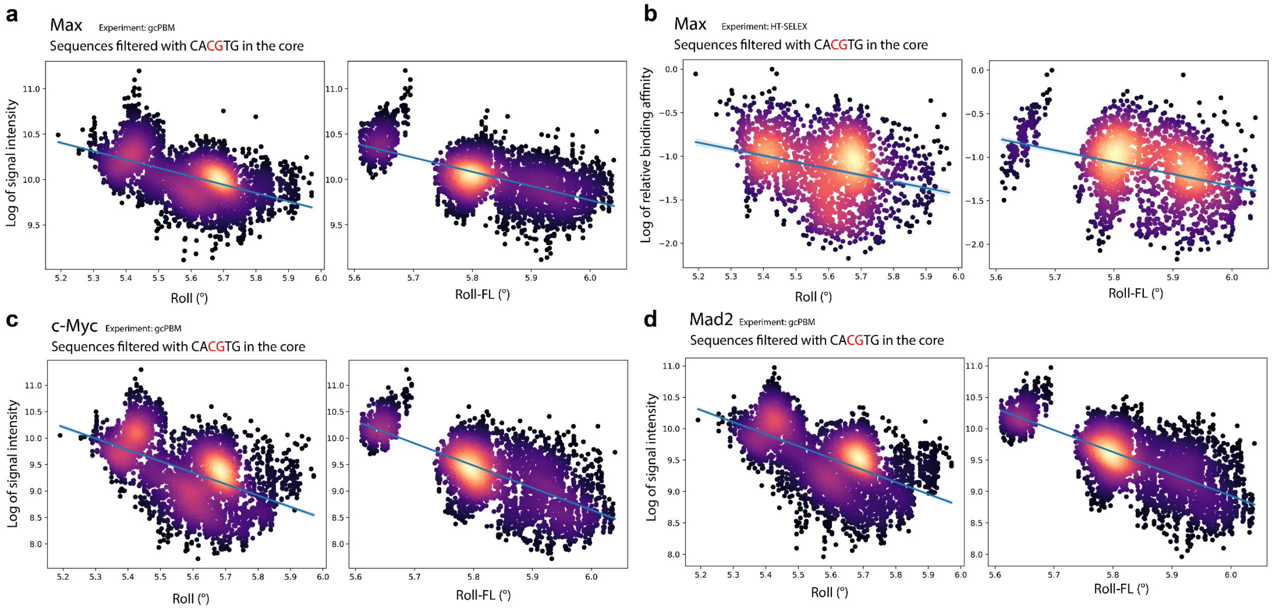


**Fig S14. Predicted Roll and Roll-FL values for Max, c-Myc and Mad2 binding data, compared to relative binding affinities from gcPBM and HT-SELEX experiment.** Data is filtered with "CACGTG" as the core region. Roll and Roll-FL features are predicted by Deep DNAshape, layer 5, for the central "CG" base pair step. Negative correlation can be found in these features.

a) Max homodimer data from gcPBM experiment.
b) Max homodimer data from HT-SELEX experiment.
c) c-Myc-Max heterodimer data from gcPBM experiment.
d) Mad2-Max heterodimer data from gcPBM experiment.

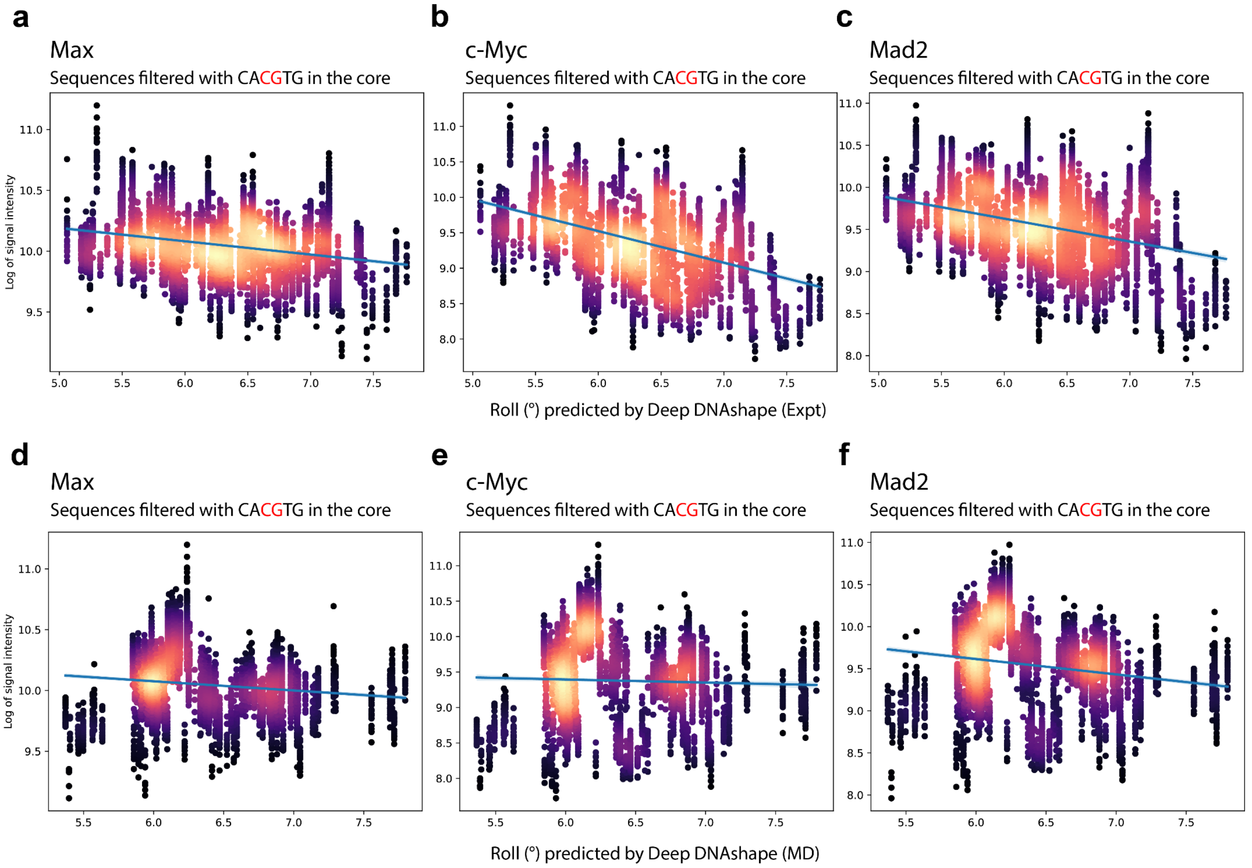


**Fig S15. Predicted Roll values for Max, c-Myc, and Mad2 binding data from gcPBM experiments, compared to the signal intensity (relative binding affinity in gcPBM).**

a-c) Roll shape features are predicted by Deep DNAshape (Expt), layer 4. Negative correlation can be found in these features.

b-f) Roll shape features are predicted by Deep DNAshape (MD), layer 4.

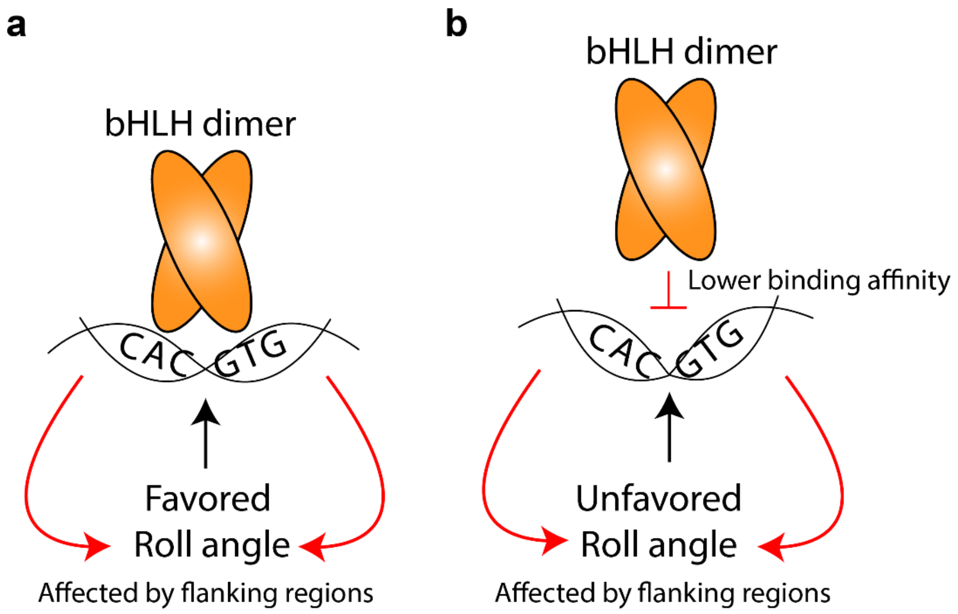


**Fig S16. Proposed binding mode of bHLH dimer using Roll readout to distinguish the same high-affinity binding sites.**

a) bHLH dimer binds to high-affinity binding sites “CACGTG”, where Roll values in the core (affected by the flanking regions) are favored.

b) bHLH dimer binds to the same high-affinity binding sites “CACGTG” with lowered binding affinity, where Roll values in the core (affected by the flanking regions) are less favored.

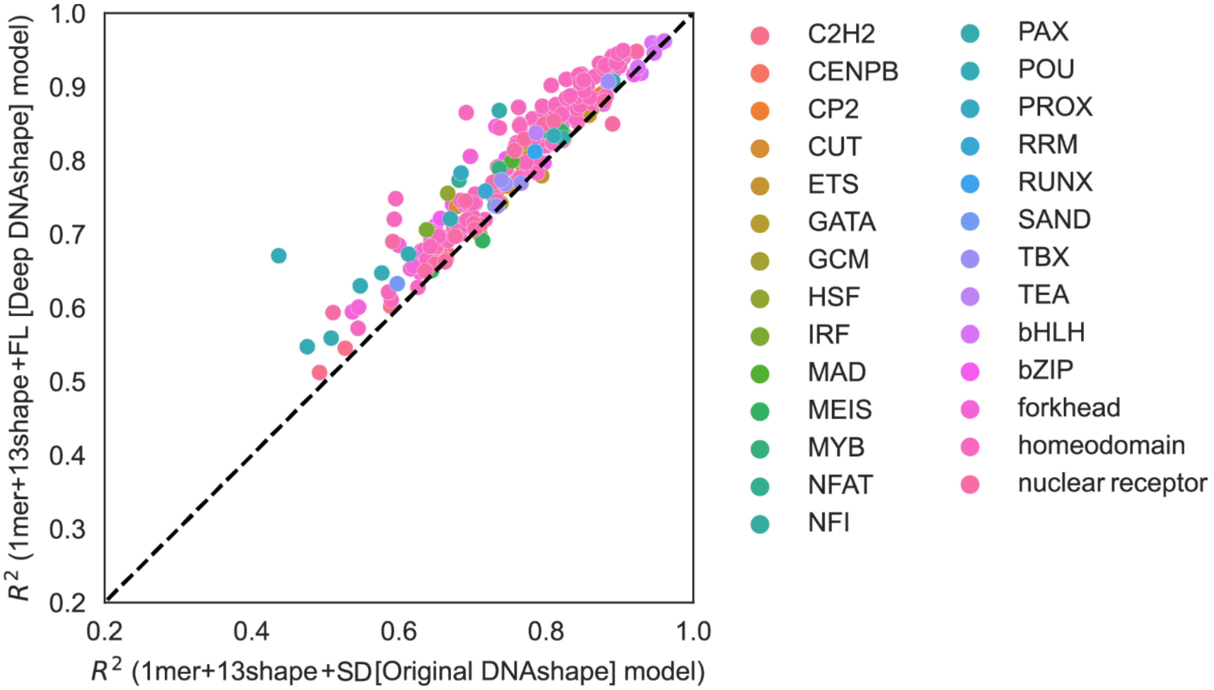


**Fig S17. $R^2$ comparison between 1mer+13shape+SD (original pentamer DNAshape) and 1mer+13shape+FL (Deep DNAshape).** SD values from original pentamer DNAshape are calculated based on statistics when compiling the raw values in the simulation data. FL values predicted by Deep DNAshape are the real shape fluctuation values encountered during simulation.

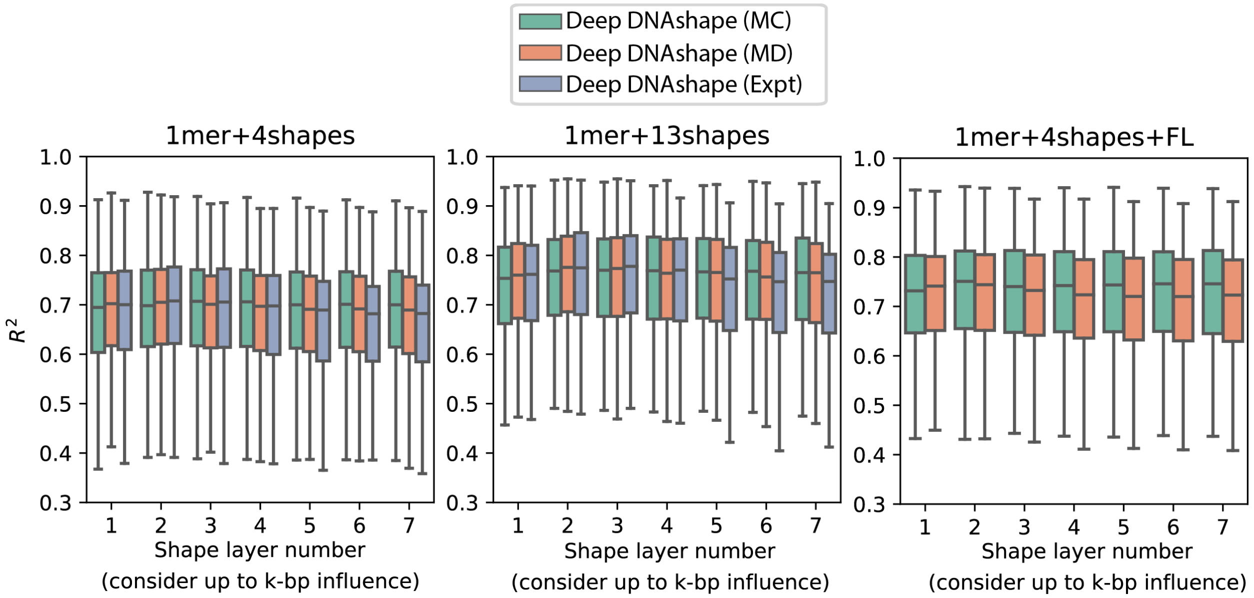


**Fig S18. $R^2$ performance of various MLR models on predicting collections of TF-DNA binding specificities from multiple experimental data.**

Shape layer number indicates which depth of shape layer is used in the Deep DNAshape models to predict DNA shape features. Significant performance differences are only found in deeper layers, where Deep DNAshape on MC data nearly always showed the best performance. Center line indicates median. Box limits are 75th and 25th percentiles. Whiskers extend 1.5 times the inter-quartile range from the top and bottom of the box. Outliers are removed in boxplots.

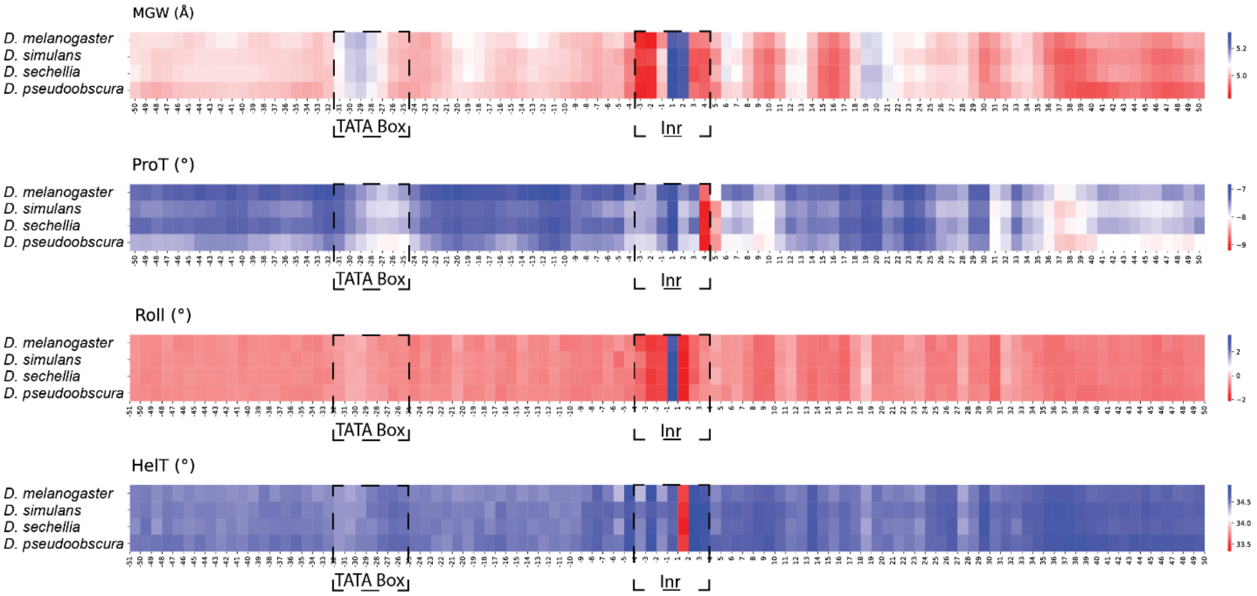


**Fig S19. Averaged DNA shape features predicted for transcription start sites (TSSs) from four *Drosophila* species[8].**

### Supplementary References


1. Young, R. T., Czapla, L., Wefers, Z. O., Cohen, B. M. & Olson, W. K. Revisiting DNA Sequence-Dependent Deformability in High-Resolution Structures: Effects of Flanking Base Pairs on Dinucleotide Morphology and Global Chain Configuration. *Life* **12**, 759 (2022).
2. Ivani, I. *et al.* Parmbsc1: a refined force field for DNA simulations. *Nat. Methods* **13**, 55–58 (2016).
3. Li, J. *et al.* Expanding the repertoire of DNA shape features for genome-scale studies of transcription factor binding. *Nucleic Acids Res.* **45**, 12877–12887 (2017).
4. Davey, C. A., Sargent, D. F., Luger, K., Maeder, A. W. & Richmond, T. J. Solvent Mediated Interactions in the Structure of the Nucleosome Core Particle at 1.9Å Resolution† † We dedicate this paper to the memory of Max Perutz who was particularly inspirational and supportive to T.J.R. in the early stages of this study. *J. Mol. Biol.* **319**, 1097–1113 (2002).
5. Lu, X. J. & Olson, W. K. 3DNA: a software package for the analysis, rebuilding and visualization of three-dimensional nucleic acid structures. *Nucleic Acids Res.* **31**, 5108–5121 (2003).
6. Zhou, T. *et al.* DNAshape: a method for the high-throughput prediction of DNA structural features on a genomic scale. *Nucleic Acids Res.* **41**, W56-W62 (2013).
7. Pasi, M. *et al.* μABC: a systematic microsecond molecular dynamics study of tetranucleotide sequence effects in B-DNA. *Nucleic Acids Res.* **42**, 12272–12283 (2014).
8. Chiu, T.-P. *et al.* GBshape: a genome browser database for DNA shape annotations. *Nucleic Acids Res.* **43**, D103-D109 (2015).
